# Transgressive segregation for salt tolerance in rice due to physiological coupling and uncoupling and genetic network rewiring

**DOI:** 10.1101/2020.06.25.171603

**Authors:** Isaiah C.M. Pabuayon, Ai Kitazumi, Kevin R. Cushman, Rakesh Kumar Singh, Glenn B. Gregorio, Balpreet Dhatt, Masoud Zabet-Moghaddam, Harkamal Walia, Benildo G. de los Reyes

## Abstract

Transgressive segregation is common in plant breeding populations, where a small minority of recombinants are outliers relative to parental phenotypes. While this phenomenon has been attributed to complementation and epistatic effects, the physiological, biochemical, and molecular bases have not been fully illuminated. By systems-level scrutiny of the IR29 x Pokkali recombinant inbred population of rice, we addressed the hypothesis that novel salt tolerance phenotypes are created by positive or negative coupling or uncoupling effects and novel regulatory networks. Hyperspectral profiling distinguished the transgressive individuals in terms of stress penalty to growth. Non-parental network signatures that led to either optimal or non-optimal integration of developmental with stress-related mechanisms were evident at the macro-physiological, biochemical, metabolic, and transcriptomic levels. The large positive net gain in super-tolerant progeny was due to ideal complementation of beneficial traits, while shedding antagonistic traits. Super-sensitivity was explained by the stacking of multiple antagonistic traits and loss of major beneficial traits. The mechanisms elucidated in this study are consistent with the Omnigenic Theory, emphasizing the synergy or lack thereof between core and peripheral components. This study supports a breeding paradigm based on genomic modeling to create the novel adaptive phenotypes for the crops of the 21^st^ century.

## Introduction

Plant breeding during the Green Revolution of the 1960’s successfully created the modern crop cultivars with superior yields under well managed environments with ideal water and nutrient conditions. Because of climate change and depletion of natural resources, the new challenges to the 21^st^ century plant breeding have become even grander (Khush, 2001, 2005; Pingali, 2012). Aided by high-throughput genotyping, marker-assisted selection, transgenic technologies, and by the emerging paradigms of gene editing and genomic selection, plant breeding has entered a new era that calls for the ability to further fine-tune genotype x environment (G x E) interaction to create novel phenotypes that were not fully addressed by the earlier breeding paradigms (de los Reyes, 2019).

Developing the next generation of crop cultivars with minimal penalty to yield under marginalized environments has become the new overarching goal of plant breeding. Key to this goal is the creation of new genetic variants and novel phenotypes. To create such novel phenotypes, a non-reductionist approach is important in considering several fundamental questions. Has plant breeding fully exhausted the potential of exotic germplasm to create novel phenotypes that are more relevant today than prior to the green revolution? With genomic biology, there is a strong optimism that precise utilization of the combining potentials of exotic germplasm could lead to further improvements in complex ‘*omnigenic*’ traits that define yield potential under marginal environments (Boyle et al., 2017). Are further genetic gains on top of what was achieved during the green revolution still possible? If so, how can the power of genomic biology be used for additional genetic gains? Genomics-enabled selection has allowed breeders to genotype each individual in breeding populations and evaluate genetic gains through the inheritance of quantitative trait loci (QTL; Li et al., 2005; Lippman et al., 2007; Thomson et al., 2010). The paradigm of *QTL stacking* promises to optimize QTL combinations to exploit the additive and non-additive potentials of both parents for maximal gains (deVicente and Tanksley, 1993; Eshed and Zamir, 1995; Tanksley and Nelson, 1996; Ashikari and Matsuoka, 2006; Kumar et al., 2018; Sandhu et al., 2018).

Both questions should inspire critical thinking to combine the classic phenomena in plant breeding with genomic biology towards the creation of novel phenotypes. Transgressive segregation is one good example of such phenomena. It was proposed that transgressive phenotypes in breeding populations are similar to the genetic novelties in natural population of hybrids that serve as foundation for adaptive speciation (Rieseberg et al., 1996; Dickinson et al., 2003; Rieseberg et al., 2003; de los Reyes, 2019). Transgressive segregation is characterized by heritable variation across the progenies of genetically divergent parents where a small minority are beyond the parental range and thus differs explicitly from heterosis (Bomblies and Weigel, 2007; Lippman and Zamir, 2007; Birchler et al., 2010). Inspired by evolutionary biology, a classic phenomenon such as transgressive segregation should be re-envisioned as a possible means to achieve further genetic gains in adaptive potentials for the 21^st^ century crop varieties.

Epistatic interactions of parental alleles and complementation effects of additive alleles are considered to be the major causes of superior or inferior attributes of transgressive segregants (Dittrich-Reed and Fitzpatrick, 2013). The recently proposed *Omnigenic Theory* defines quantitative traits in terms of both additive and non-additive contributions of few large-effect core loci and hundreds of minute-effect peripheral loci across the genome (Boyle et al., 2017). We further hypothesized that the superiority or inferiority of transgressive segregants are due to ideal coupling/uncoupling or non-ideal coupling/uncoupling, respectively, of the various compatible and incompatible biochemical and developmental attributes from each parent encoded by the core and peripheral loci. Such mechanisms lead to either physiological gain or physiological drag in certain individuals, determining positive or negative net gain (de los Reyes, 2019). Because of the many potential attributes that should be affected by such coupling and uncoupling mechanisms, it would take many generations of genetic recombination and genome reshuffling to observe the physiological gain or drags on a relatively small number of individuals in a segregating population.

To test the physiological coupling and uncoupling and network rewiring hypotheses, we employed a multi-tier approach to examine a well characterized recombinant inbred population of rice for salt tolerance that was developed by the International Rice Research Institute (IRRI) in the 1990’s from the parental cultivars IR29 (*Xian/Indica*; salt-sensitive) and Pokkali (*Aus*; salt-tolerant) (Bonilla et al., 2002; Gregorio et al., 2002; Walia et al., 2005; Singh et al., 2007; Thomson et al., 2010). Pokkali is a photoperiod-sensitive landrace historically used as donor of salt tolerance in rice breeding programs in India (Moeljopawiro and Ikehashi, 1981; Sahi et al., 2006). We were particularly interested in exploring the full combining potential of this donor with a salt-sensitive but high-yielding *Xian/Indica* cultivar (IR29) beyond what has been shown by QTL mapping. QTL mapping conducted on this population identified the *Saltol* on chromosome-1 of Pokkali to be responsible for much of phenotypic variance for salt stress defenses at the seedling stage. *Saltol* included genes involved in the regulation of Na^+^ uptake and accumulation in the shoots (Ren et al., 2005; Thomson et al., 2010). The recombinant inbred line (RIL, F_8_) FL478 has served as the non-photoperiod sensitive donor of *Saltol* to improve the seedling-stage salt tolerance of *Xian/Indica* rice cultivars (Huyen et al., 2012; Islam et al., 2012; Huyen et al., 2013; Chattopadhyay et al., 2014; Bimpong et al., 2016; Waziri et al., 2016; Babu et al., 2017).

We describe here the results of a multi-tier macro-physiological, biochemical, and molecular profiling of the IR29 x Pokkali RILs at the population and individual levels. We scrutinized specific individuals representing each phenotypic class by integrating multi-level ‘omics’ datasets. With this approach, we were able to identify clear transgressive segregants and provide strong support to the coupling and uncoupling hypothesis by revealing the hidden potentials of the salt-sensitive *Xian/Indica* parent IR29 towards positive complementation with *Saltol* and other genes from Pokkali. We also revealed the potential of Pokkali as a source of physiological drags against other minor effect genes. This study also provides direct evidence that transgressive traits are created by network rewiring as an outcome of recombination.

## Results

### Evaluation of the IR29 x Pokkali RILs under severe salt stress

During the fine-mapping of *Saltol*, the IR29 x Pokkali *core mapping population* (F_2_, F_8_-RILs of 126 individuals) was evaluated at IRRI at the seedling stage (V1 to V5) under EC = 9 dS/m for Na^+^ exclusion potential, with shoot and root Na^+^/K^+^ and Standard Evaluation Score (SES) as main criteria (Counce et al., 2000). The SES ranges from 1 to 9, with 1 representing the highest level of tolerance. IRRI selected 64 F_8_-RILs (*representative group*) spanning the range of salt tolerance across the population. Along with the parents IR29 and Pokkali, and an additional salt-sensitive check IR64, we conducted a time-course (0, 24, 48, 72, 144 hr) evaluation of salt tolerance across the *representative groups* at tillering stage (V4 to V12). A more severe salt stress (EC = 12 dS/m) was used to further push the limits of phenotypic potentials that were not revealed by the earlier milder treatments. Using SES as initial criterion, we expected to identify outlier RILs that perform worse than IR29 or better than Pokkali at V4 to V12. Results revealed slightly different ranking with clear outliers not identified at V1 to V5 (Supplemental Data Set 1). From these, we established the *minimal comparative panel*, consisting of thirty seven (37) RILs spanning the full range of tolerance revealed at EC = 9 dS/m (sensitive = IR29, FL454; tolerant = Pokkali, FL478), and the outliers revealed at EC = 12 dS/m, consisting of the super-sensitive FL499 and super-tolerant FL510 (Table 1, Figure 1A).

**Table 1.** Summary of the Standard Evaluation Scores (SES) and qualitative phenotypic ranking at tillering stage (V4 to V8) across the 67 F_8_-RILs (*i.e*., representative group; *n = 8*).

| Genotype | SES<br>EC=12<br>(TTU) | Classification | Saltol Peak<br>Marker (+/-) | Saltol Flanking<br>markers (+/-) | Genotype | SES<br>EC=12<br>(TTU) | Classification | Saltol Peak<br>Marker (+/-) | Saltol Flanking<br>markers (+/-) |
| --- | --- | --- | --- | --- | --- | --- | --- | --- | --- |
| FL301 | 4.5 |  | + | - | FL460 | 5.5 |  | - | - |
| FL302 | 4 | Moderate | + | NA | FL463 | 5 |  | - | - |
| FL303 | 4.5 | Sensitive | - | NA | FL468 | 7 |  | - | - |
| FL327 | 5.5 | Sensitive | + | + | FL469 | 3.5 | Moderate | - | - |
| FL333 | 4.5 | Sensitive | + | + | FL472 | 7 |  | NA | + |
| FL352 | 5 | Sensitive | - | - | FL473 | 4.5 |  | + | + |
| FL356 | 7 |  | - | - | FL476 | 7 | Sensitive | + | NA |
| FL358 | 7 | Sensitive | + | NA | FL478 | 3 | Moderate | + | - |
| FL378 | 2.5 | Tolerant | + | + | FL483 | 4.5 |  | - | NA |
| FL382 | 3.5 | Moderate | - | NA | FL494 | 3 | Moderate | - | - |
| FL387 | 4.5 | Moderate | + | + | FL496 | 2 | Tolerant | + | NA |
| FL389 | 6 |  | - | - | FL499 | 9 | Supersensitive | NA | + |
| FL395 | 6 | Sensitive | - | - | FL501 | 2 | Tolerant | + | + |
| FL396 | 3.5 | Moderate | + | NA | FL502 | 3 | Moderate | - | - |
| FL397 | 4 |  | - | + | FL507 | 7 |  | + | + |
| FL399 | 6 |  | - | NA | FL508 | 6 |  | + | NA |
| FL407 | 2 | Tolerant | + | + | FL510 | 1.5 | Supertolerant | + | - |
| FL409 | 6 |  | - | - | FL512 | 8 |  | - | - |
| FL416 | 2.5 | Tolerant | NA | + | FL530 | 3 | Moderate | + | + |
| FL418 | 7 |  | - | - | FL541 | 4 |  | + | + |
| FL428 | 2 | Tolerant | + | - | FL542 | 7 |  | + | + |
| FL431 | 4.5 | Sensitive | - | - | FL545 | 4 | Moderate | + | - |
| FL433 | 4 |  | - | - | FL557 | 3.5 | Moderate | - | - |
| FL434 | 4.5 |  | - | NA | FL560 | 2.5 | Tolerant | + | + |
| FL435 | 5 |  | - | - | FL561 | 6 | Sensitive | + | + |
| FL441 | 7 |  | - | + | FL567 | 4 | Moderate | - | - |
| FL442 | 7 |  | + | - | FL568 | 4 | Moderate | - | NA |
| FL443 | 7 |  | NA | - | FL572 | 6 |  | - | - |
| FL445 | 7 |  | + | + | FL577 | 6 |  | + | NA |
| FL446 | 5 |  | - | + | FL588 | 6 |  | + | NA |
| FL448 | 4 | Moderate | - | - | FL590 | 6 |  | + | NA |
| FL449 | 2.5 | Tolerant | + | - | IR29 | 8 | Parent (Sensitive) | - | - |
| FL454 | 4.5 | Sensitive | - | - | IR64 | 3.5/8 | Check (Moderate) | NA | NA |
| FL456 | 7.5 |  | NA | + | Pokkali | 5 | Parent (Moderate) | + | + |

Salt tolerance was compared with *Saltol* genotype, which was determined based on the presence of the Pokkali allele at the peak and/or flanking markers (Thomson et al., 2010).

**Figure 1.**
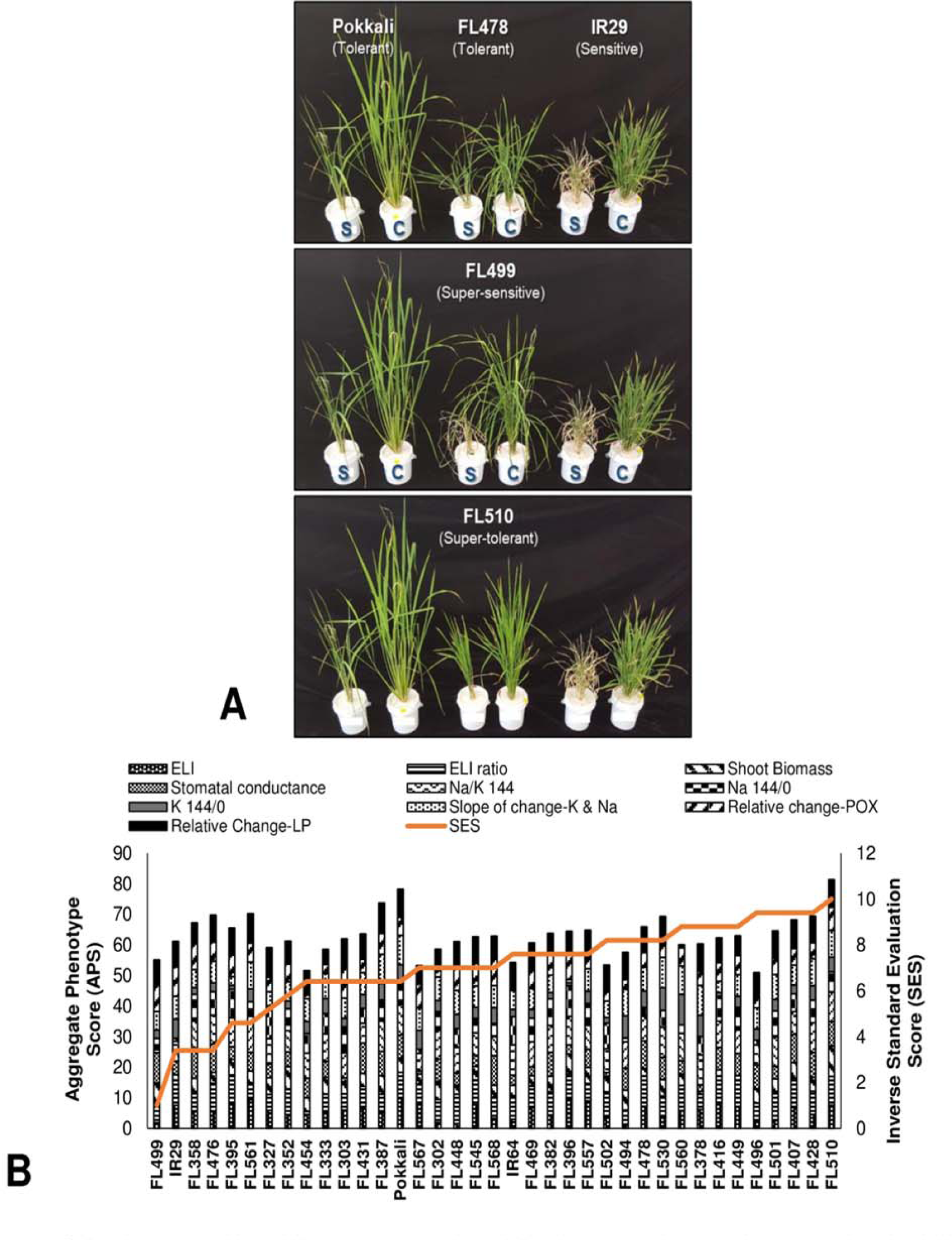
Salt tolerance at V4 to V8 stage across the *minimal comparative panal* representing the full phenotypic range at EC=9 dS/m (IRRI) and EC= 12 dS/m(Texas). **(A)** Comparison of health in control (C) and stress (S) experiments after seven days (144 hrs) at EC= 12 dS/m The outliers FL510 (super-tolerant) and FL499 (super-senstive) are highlighted compared to tolerant parent Pokkali (*Saltol* donor), sensitive parent IR29, and tolerant sibling FL478 (*see Table 1*). Differences in injury and growth were evident particularly between the transgressive FL510 and FL499. **(B)** individual physio-morphometric scores were normalized and combined as *Aggregate Phenotypic Score* (APS), which assumes equal weights of each parameter that includes Electrolyte Leakage index (ELI) and its ratio (ELI ratio) at first injury (72 hr) and maximum injury (144 hr), shoot biomass ratio (stress/control), stomatal conductance ratio (stress/control), Na^+^/K^+^ at maximum injury (Na/K 144/0),Na^+^ ratio at maximum injury (Na 144/0), K^+^ ratio at maximum injury (K 144/0), slope of K^+^ and Na^+^ changes (control/stress), change in peroxidase activity (change-POX; stress/control), and change in lipid peroxidation (change-LP; stress/control). APS was plotted against the inverse of SES for direct proportionality.

The *minimal comparative panel* was further profiled for a set of physio-morphometric parameters including electrolyte leakage index (ELI), cellular ion concentration, lipid peroxidation (LP) as a measure of cellular membrane injury, total peroxidase activity (POX) as measure of ROS scavenging capacity, shoot biomass, and stomatal conductance (Figure 1B). Only mild correlation with SES was established (Supplemental Table 1), including POX at the point (144 hr) of maximum injury (r^2^ = 0.17), shoot Na^+^/K^+^ (r^2^ = 0.15), and POX stress/control ratio (r^2^ = 0.11). Collectively, the physio-morphometric parameters did not correlate strongly with SES.

The tolerant FL478 carrying an introgression for *Saltol* allele of Pokkali was as tolerant as Pokkali under both EC = 9 dS/m and EC = 12 dS/m (Table 1, Figure 1A; (Thomson et al., 2010). Other RILs with homozygous introgression for Pokkali *Saltol* allele (Supplemental Data Set 1) also had comparable SES as Pokkali and FL478, confirming the biological significance of the blind SES ranking at EC = 12 dS/m. As an outlier, the super-tolerant FL510 had the best SES and highest aggregate phenotypic score (APS) across the population, significantly outperforming the tolerant Pokkali and FL478. While the individual scores of FL510 were not always the best, none of them were below the population means, implying the additive effects of many minutely contributing parameters to its overall potential. FL478 tended to have very good scores in some parameters but poor scores in others, which appeared to cause a penalty to the net phenotypic score. Pokkali had a very good APS, but its SES was inferior to FL510 (Figure 1B).

The sensitive parent IR29 along with about 30% of RILs including FL454 (sensitive) and FL499 (super-sensitive) were the worst performers based on SES alone (Supplemental Data Set 1; Table 1). FL499 was severely injured with the worst scores for most parameters especially for ELI and Na^+^/K^+^. The trends in APS showed that the poor ranking of FL499 was due to poor scores in the majority but not all parameters (Figure 1B). Integration of all physio-morphometric parameters into an aggregate score (APS) indicate the various permutations by which different parental attributes could be combined in the progenies either optimally or non-optimally.

### Relatedness among RILs based on physio-morphometric matrix

Based on few parameters where the sensitive parent IR29 had significantly better scores than the tolerant parent Pokkali, we hypothesized that some of IR29-derived properties may be contributing to tolerance potential when combined with other complementary properties from Pokkali. To address this hypothesis, we used the total matrix to measure the similarities across RILs through neighbor-joining dendrograms. The first measure established global relationships based on all contributing factors. The second was based only on Na^+^ exclusion components (Na^+^ content, K^+^ content, Na^+^/K^+^, ELI) to assess the contribution of *Saltol* (Table 1).

The full-matrix dendrogram illustrates the overall range of similarities (Figure 2A), where the super-sensitive FL499 formed the most distant clade. The other poor RILs (*e.g*., IR29, FL454) formed distinct clades from the better RILs (*e.g*., Pokkali, FL478, FL510). The super-tolerant FL510 also formed the most distant sub-clade among the mostly good RILs, consistent with its superior tolerance. However, the large clade of mostly good RILs also included few inferior RILs, suggesting that even the poor RILs shared some attributes with their better siblings.

**Figure 2.**
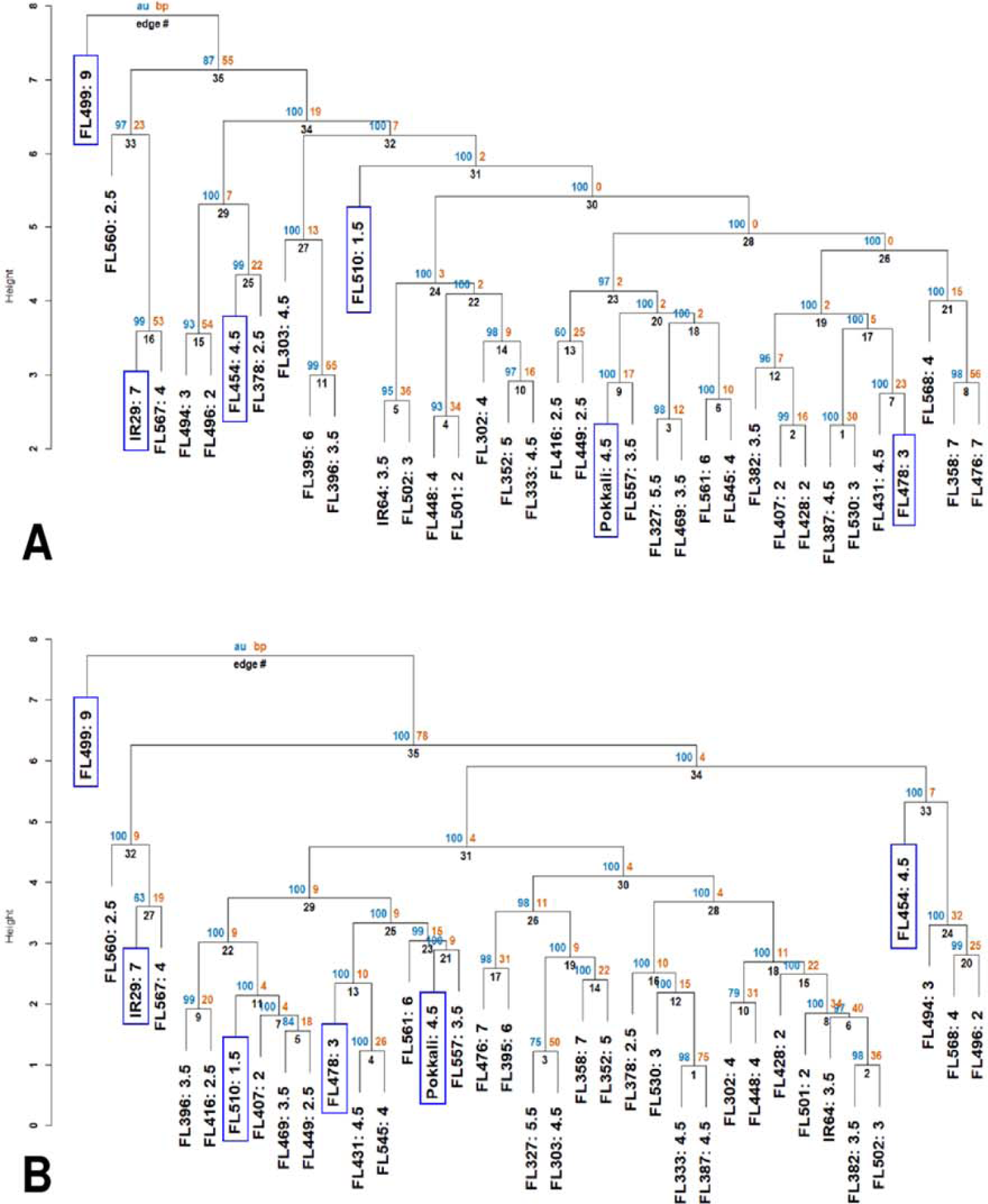
Neighbor-joining dendrograms showing two layers of similarities based on the phenotypic matrix (APS, SES). Each genotype in the dendrogram is suffixed with their SES. The individual genotypes (parents, RILs) that were investigated to understand the physiological mechanisms are highlighted in blue boxes. AU (blue) refers to the adjusted P-value of clustering, and BP (orange) refers to the bootstrapping value of ‘pvclust’ package. **(A)** Similarities based on the entire matrix (components of APS + SES). Dendrogram shows the poor genotypes represented by IR29, FL454, and FL499 forming distinct clades from the good genotypes represented by Pokkali, FL478, and FL510. **(B)** Similarities based on APS components relevant to Na^+^ sequestration (ELI, Na^+^ and K^+^ contents at control and 144 hr, Na^+^/K^+^ ratio). Dendrogram shows the tendency for individuals carrying the Pokkali *Saltol* allele to cluster together within one large clade. In both **(A)** and **(B)**, the super-sensitive FL499 had the earliest divergence.

*Saltol* effects are primarily associated with Na^+^ exclusion. The dendrogram based on salt exclusion components revealed even more meaningful groupings (Figure 2B). The clear outliers were the inferior RILs (*e.g*., IR29, FL454, FL499) that are negative for *Saltol* (Table 1). The large clade consisting of mostly good RILs is divided into two sub-clades. The first is comprised of RILs that are positive for *Saltol* (tolerant Pokkali and FL478, and super-tolerant FL510), with Pokkali and FL478 being more closely related to each other than to FL510. The second is comprised of mostly (13 out of 17) good and moderate RILs that are positive for *Saltol*, indicating that the similarities among the good RILs are largely due to *Saltol*. However, such similarities excluded the contributions of other properties in the full matrix, hence without the contributions of other factors in the genetic background.

### Real-time profiling of growth and physiological potentials

We used the representative phenotypic classes comprised of the sensitive parent (IR29), tolerant parent (Pokkali), sensitive RIL (FL454), tolerant RIL (FL478), super-sensitive RIL (FL499), and super-tolerant RIL (FL510) to track plant growth in real-time with the LemnaTec imaging system for a period of 18 days from V4 to V12. RGB imaging was used to determine the projected shoot area (PSA) and plant height, which are correlated with plant biomass (Campbell et al., 2015). Plants were stressed at EC = 9 dS/m to remove potential bias against the sensitive classes given that EC = 12 dS/m was nearly lethal for FL499. Nevertheless, EC = 9 dS/m was strong enough to differentiate the potentials across the panel.

The image-based growth curves based on daily measurements of plant area and height showed linear growth patterns over time (Figure 3A). However, major differences even in the control were evident in terms of inherent growth potentials. Growth rates in Pokkali, FL454, and FL499 were much higher compared to IR29, FL478 and FL510 even in the control. In the RILs, the growth potentials appeared to be due to the combined effects of parental attributes. For instance, the fast-growing salt-sensitive RILs (FL454, FL499) were more like the salt-tolerant parent Pokkali in terms of plant height but their tillering capacity was more similar to IR29. The salt-tolerant RILs (FL478, FL510) were more similar in overall growth pattern to the salt-sensitive parent IR29. The high-yielding improved *Xian/Indica* variety IR29 has short stature with profuse tillering, while the *Aus* landrace Pokkali has tall stature, darker green leaves, and less tillering. FL478 has the same *Xian/Indica* plant-type as IR29, while FL510 has distinct morphology (Figure 1A). It is intermediate in height, with thick, dark green, and erect leaves and stems, similar to the new plant-type (NPT) bred at IRRI during the 1990’s (Peng et al., 1994; Peng et al., 2008).

**Figure 3.**
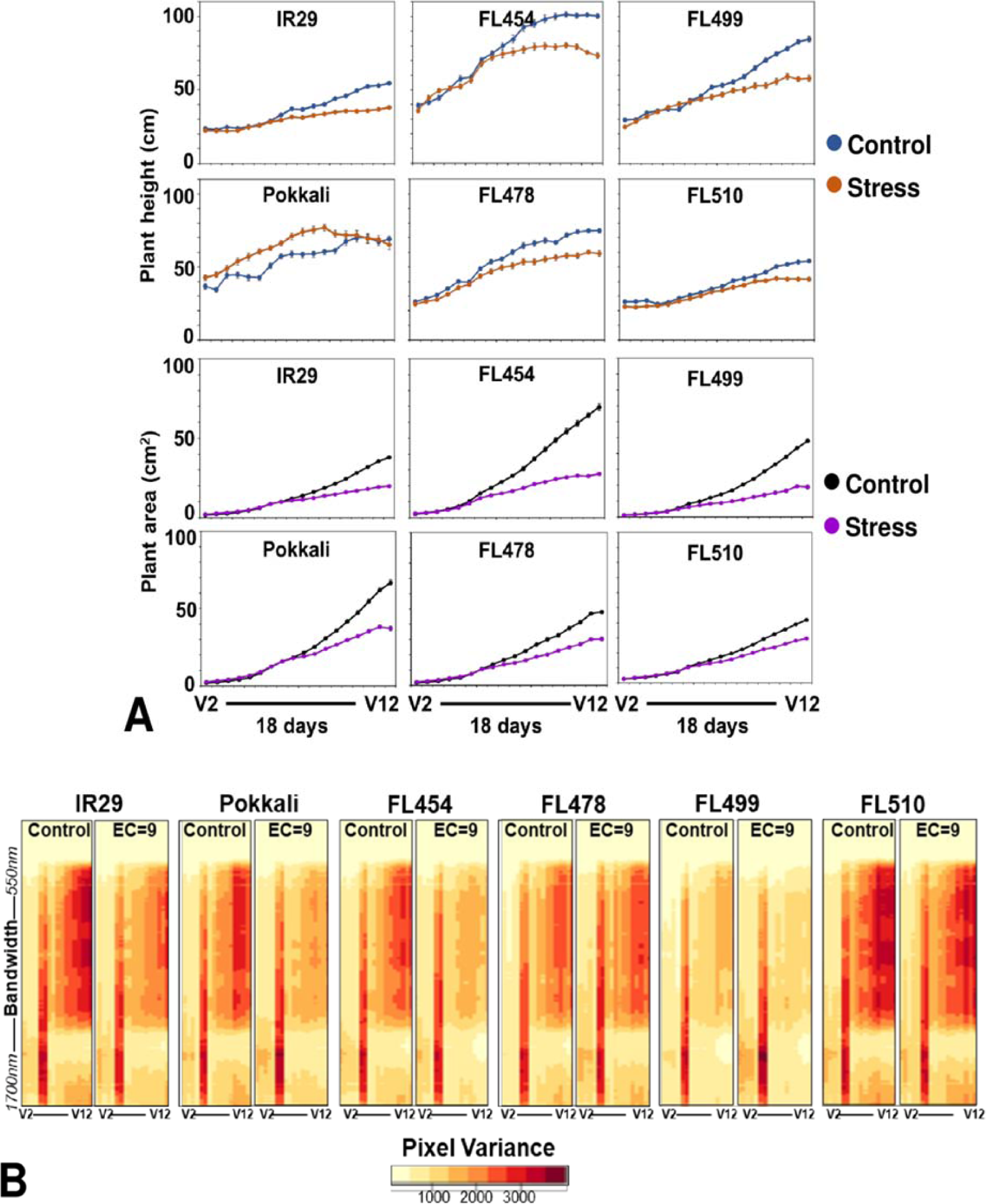
Real-time growth profiling of the sensitive parent IR29, tolerant parent Pokkali, sensitive RIL FL454, tolerant RIL FL478, super-sensitive RIL FL499, and super-tolerant RIL FL510 at EC= 9 dS/m. **(A)** Growth curves as a function of changes in plant area and plant height during the 18-day period of stress, calculated based on pixels captured by RGB and hyperspectral camera. Note the time of forking between the control and stress curves and the angle of the fork, reflective of growth plateauing. **(B)** Variation in hyperspectral variance at 243 wavelength bands ranging from 550nm to 1700nm. Heat maps of hyperspectral variances are potential indicators of overall plant health and stress injuries. The overall patterns in hyperspectral variances mirror the patterns revealed by the growth curve analysis.

Growth curves indicate that stress penalty was common across all genotypes (Figure 3A). However, the magnitude of penalty varied significantly regardless of the inherent variation in growth potentials under control condition. For instance, the timing when growth retardation occurred varied with clear correlation to salt tolerance. Plants under control and stress grew at similar rates before stress. While growth retardation based on projected shoot area (PSA) and height was detectable as early as seven (7) days after stress, variation across genotypes was evident based on the timing of the forking between the control and stress curves, which determined the magnitude of growth plateauing (*i.e*., fork angle) in real-time (Figure 3A). Growth penalty was most severe in the salt-sensitive IR29, FL454 and FL499 relative to their respective controls. This was also supported by few other parameters such as tiller reduction, plant biomass, and symptoms of leaf injury. Tillering of Pokkali and FL478 were also negatively affected but in much lesser magnitude than IR29, FL454, and FL499. The super-tolerant FL510 had the least penalty based on the degree of tiller reduction and timing and angle of forking (Figure 3A). The PSA ratio (stress/control) also indicate difference in tolerance based on growth maintenance (Supplemental Figure 1). Tiller reduction was evident from decreasing PSA ratio in all genotypes. However, FL510 and FL478 stayed close to a ratio of 1, indicating much less damage.

To further substantiate the observed variation in growth penalty, we examined the hyperspectral profiles across the growth window as potential indicator of plant health and vigor. Hyperspectral profiles were based on image captured at 243 wavelength bands ranging from 550nm to 1700nm, with each band evenly separated by 4.7 nm. Image variance plots described the spread of pixel intensities as indicators of how much physical changes including wilting, drying, yellowing or necrosis occurred during stress.

Differences in growth across genotypes (Figure 3A) were generally consistent with the trends in hyperspectral profiles (Figure 3B). Reduction in image variances was quite apparent in IR29 and FL454 but only minimal in FL510 and FL478, indicative of lesser injuries. These genotypes had more stable profiles throughout the stress period. FL499 had lower image variances even under control, suggestive of less than optimal growth that was further exacerbated by stress. Interestingly, the hyperspectral profiles of the super-tolerant FL510 in both control and stress were quite similar to the high-yielding parent IR29 under control. This suggests a positive attribute acquired by FL510 from IR29, which translated into net gains in combination with other defense-related traits inherited from Pokkali.

### Na^+^/K^+^ and proline profiles in relation to real-time growth variances

The dynamics of Na^+^ and K^+^ accumulation was investigated to assess how much the capacity for Na^+^ exclusion contributed to the growth penalty in sensitive and tolerant genotypes. The Na^+^/K^+^ ratio (Na^+^/K^+^) in shoots and roots were used as primary indicators of tolerance. Variation for Na^+^/K^+^ in shoot across genotypes was significant in stress but not in control (Supplemental Figure 2A). IR29, FL454, and FL499 had significantly higher Na^+^/K^+^ than Pokkali, FL478, and FL510. The mean Na^+^/K^+^ segregated the panel into tolerant (Pokkali, FL478, FL510) and sensitive (IR29, FL454, FL499) groups (P < 0.05). Because of *Saltol*, Pokkali had the lowest Na^+^/K^+^ within the tolerant group, thus the overall potentials of FL478 and FL510 may not dependent on *Saltol* effects alone. No significant difference in root Na^+^/K^+^ was detected across genotypes (Supplemental Figure 2B).

Proline is known to contribute to the maintenance of cellular water potential by osmotic adjustment (Claussen, 2005; Ashraf and Foolad, 2007; Kumar et al., 2010; Szabados and Savoure, 2010). The total proline content of each genotype was measured after seven (7) days of salt stress, which also correspond to the initiation of the forking between the control and stress growth curves (Figure 3A). Proline content increased significantly (P < 0.05) under salt stress in IR29 and FL454 but not in other genotypes (Supplemental Figure 2C), suggesting that osmotic adjustment is not a major contributing factor to the variation in growth penalties.

### Metabolite profiles across the gradient of salt tolerance potential

Shotgun profiling by LC-MS/MS was performed to compare the stress metabolite signatures across genotypes. About 7,000 unique macromolecules were identified, many of which were unique to a given genotype. A total of 217 known metabolites had differential abundances between stress and control at P < 0.05 (Supplemental Data Set 2). K-means clustering and principal component analysis (PCA) revealed multiple components that spread the entire dataset widely based on variances. FL499 had the most distinct profile compared to the other genotypes (Figure 4A). The PCA plot revealed significant overlaps among the tolerant genotypes (FL478, FL510) for the differentially abundant metabolites, with patterns completely opposite to the profiles of the parents (IR29, Pokkali) and super-sensitive FL499. These trends indicate metabolite signatures associated with salt tolerance. The super-tolerant FL510 was more similar to the sensitive parent IR29, while the tolerant FL478 was more similar to the tolerant parent Pokkali (Figure 4B). All these suggest that the improved high-yielding but salt-sensitive parent IR29 has significant contributions to the overall potential of the transgressive super-tolerant FL510.

**Figure 4.**
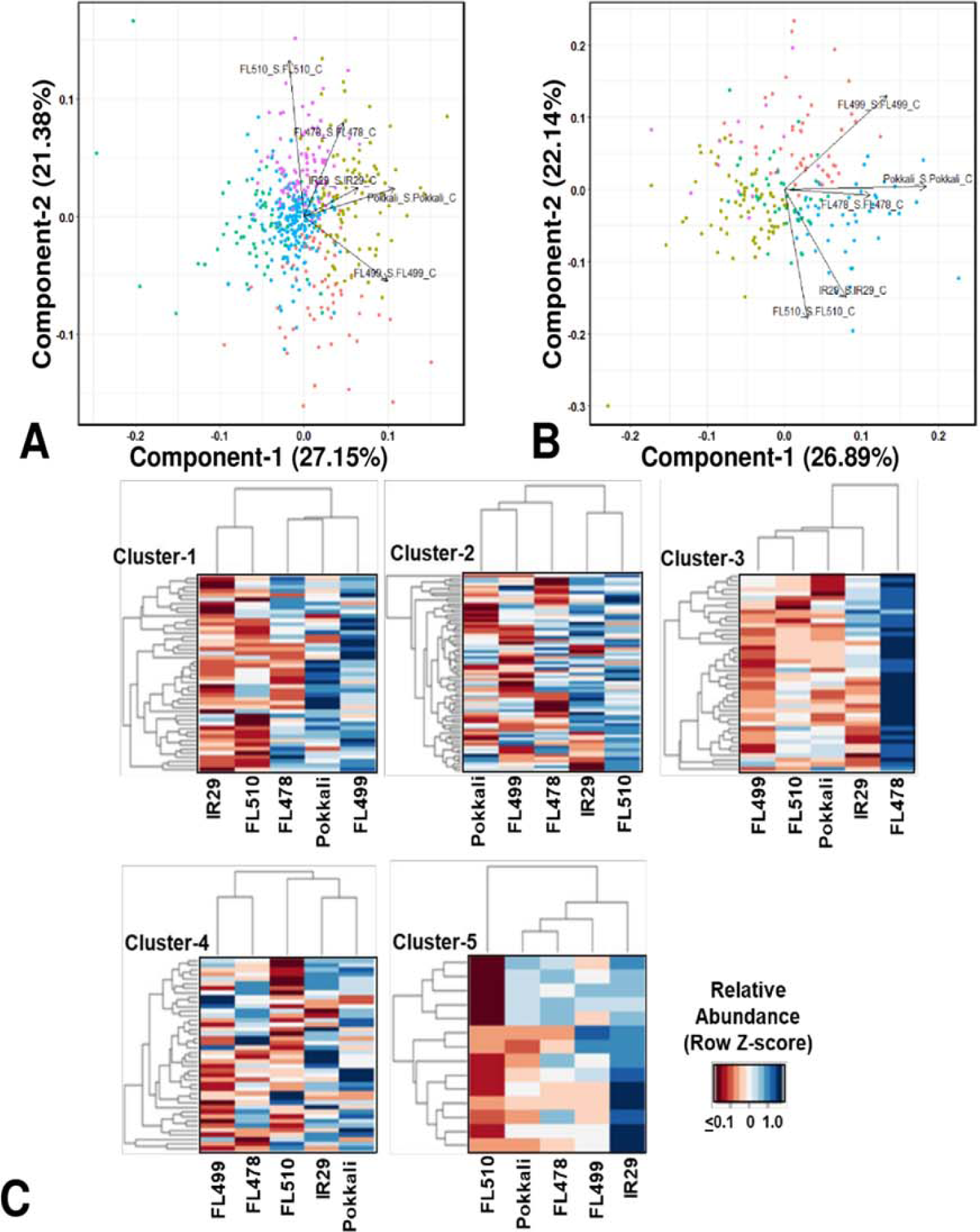
Shotgun LC-MS/MS metabolite profiles across the representative phenotypic classes at the point of maximum stress (7 days or 144 hr) at EC= 12 dS/m. **(A)** Principal component analysis (PCA) of metabolites with common occurrence regardless between control and stress in sensitive parent IR29, toierant parent Pokkali, sensitive FL454, tolerant FL478, super-sensitive FL499, and super-tolerant FL510. PC-1 and PC-2 separated the good RILs FL478 and FL510 from the poorest RIL FL499, accounting for 27.15% and 21.38% of phenotypic variance, respectively. The PCA also shows high similarities between the two parents. **(B)** PCA of metabolites with differential abundances between control and stress at p< 0.05, Based on PC-1 and PC-2, which accounted for 26.89% and 22.14% of phenotypic variance, respectively, the super-sensitive FL499 was quite distant from the super-tolerant FL510, despite the high level of similarities between the parents, hence non-additive and transgressive. The tolerant FL478 was more similar to Pokkali than to IR29, and any of its siblings. **(C)** K-means clustering heat maps and dendrograms showing the patterns of metabolite co-abundances across genotypes. Heatmap highlights a cluster that is highly abundant in each genotype. Clustering by genotype shows the similarity of IR29 and FL510, especially in clusters 1, 2, and 4. Cluster 5, highlights the differences that are contributory towards contrasting phenotypes.

The K-means clustering of differentially abundant metabolites revealed different patterns of co-expression (Figure 4C). Clusters-1, 2 and 4, which are enriched with different types of flavonoids and other antioxidants have similar profiles in IR29 and FL510 but not in other genotypes. Cluster-1 is highlighted by known indicators of oxidative stress such as (-)-allantoin (Kand’ár and Žáková, 2008) and distinguishes the super-sensitive FL499 from the others. Cluster-3 distinguished the tolerant FL478 from the rest, enriched with JA-associated metabolites and fatty acids.

ABA and other related metabolites are the dominant components of cluster-4. This cluster is most similar between the tolerant parent Pokkali, sensitive parent IR29 and tolerant FL478, and separated the super-tolerant FL510 and super-sensitive FL499 from the group. Cluster-5 features the compatible osmolyte trehalose and other compounds with similar properties and distinguished the sensitive parent IR29. Overall, metabolite profiles revealed meaningful enrichments that distinguished the transgressive segregants. PCA indicates that the salt-sensitive parent IR29 has important contributions to the overall potential of the super-tolerant FL510, consistent with the trends revealed from the real-time growth profiling.

### Transcriptome signatures across the gradient of salt tolerance

The leaf transcriptomes of each genotype were profiled in a time-course (0, 24, 48, 72,144 hr) RNA-Seq experiment to further substantiate the properties of the outliers. The RNA-Seq mapping statistics of 44,199 transcript variants from 14,696 expressed loci are summarized in Supplemental Table 2. Contrasting profiles were evident between IR29 and Pokkali, while similarities and differences were evident across RILs (Figure 5). The super-tolerant FL510 had the highest number of genes with no significant change in expression at EC = 12 dS/m. Upregulation in FL510 occurred gradually but progressively compared to others. In the super-sensitive FL499, only subtle changes occurred during the early stages of stress, but many genes were drastically upregulated or downregulated at 144hr/7-day, coinciding with severe leaf senescence and necrosis. The tolerant FL478 and sensitive FL454 had large number of early, late, sustained, and transiently upregulated and downregulated genes similar to IR29 and Pokkali.

**Figure 5.**
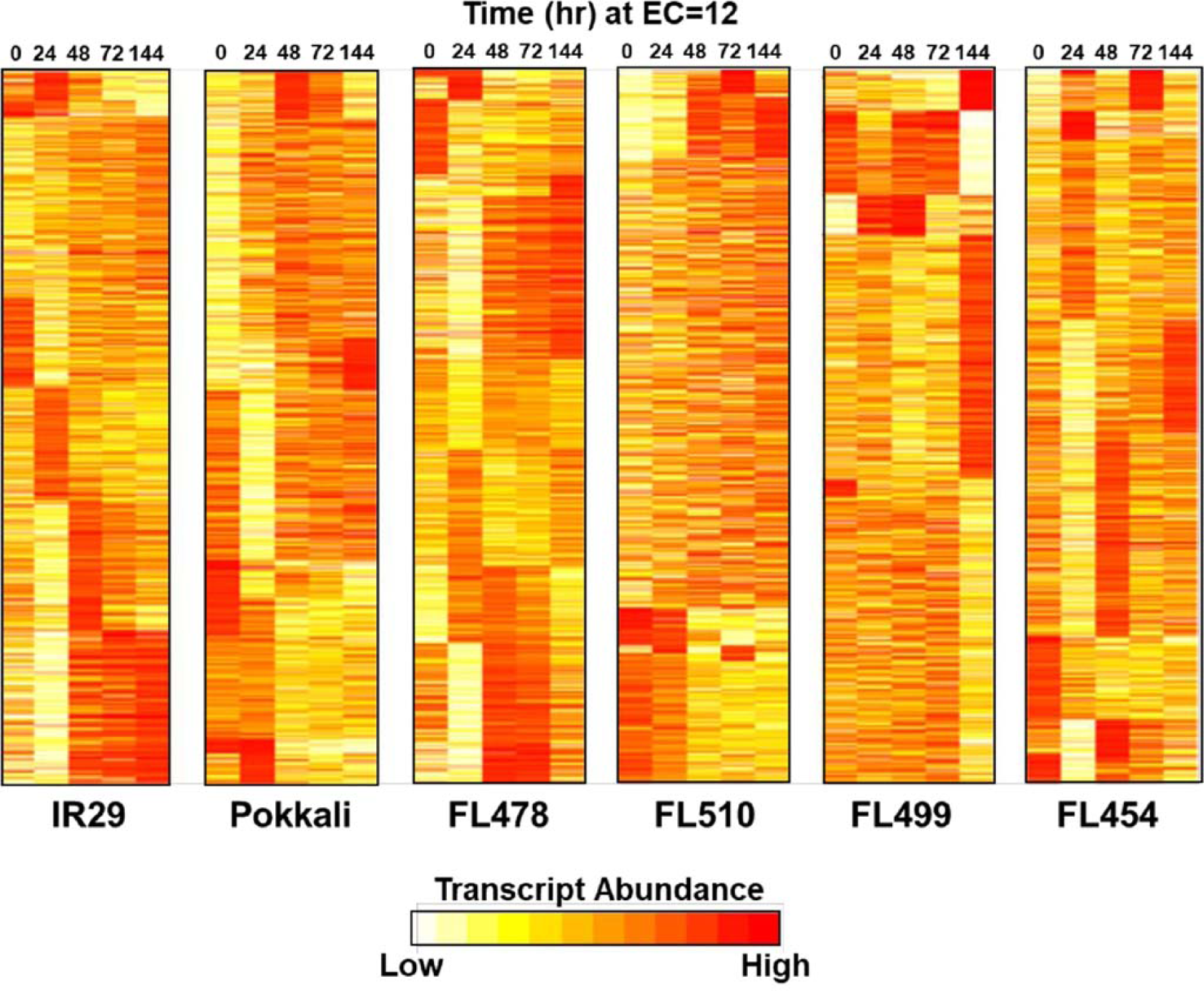
Temporal profiles of salt stress (EC= 12 dS/m) transcriptomes in parents (IR29 = sensitive; Pokkali = tolerant) and their RILs (FL454 = sensitive; FL499 = super-sensitive; FL478 = tolerant; FL510 = super-tolerant). Parallel comparison of transcriptomes across genotype shown as K-means++ clusters included 14,696 unique transcript loci. The super-tolerant FL510 is the most unique, having gradual changes in expression. The Othergenotypes have large clusters of co-expressed genes with drastic upregulation or downregulation across time.

The most significant changes in expression involved genes with important roles in integrating stress and growth responses, including calmodulin-like *OsCML27* (Os03g0331700) that functions as a sensor for Ca^2+^-mediated stress signaling (Perochon et al., 2011), high-affinity K^+^ transporter *OsHKT7* (Os04g0607600), which is a strongly selective transporter of Na^+^ against K^+^ (Suzuki et al., 2016; Oda et al., 2018), metallothionein *OsMTI4A* (Os12g0570700) involved in oxidative defenses (Zhou et al., 2006; Yang et al., 2009; Kumar et al., 2012), and the Multi-Pass MYB-transcription factor *OsMPS* (Os02g0618400) involved in growth regulation (Schmidt et al., 2013). These genes were used to bait for other co-expressed genes (network cohorts) to understand the significance of co-expression signatures across the comparative panel.

The *OsCML27* profile was most distinct in the super-tolerant FL510 where it was upregulated in sustained manner (Figure 6A). The parents IR29 and Pokkali did not share any cohort genes with any progeny, while only minimal overlaps at best occurred between FL510, FL478, FL499, and FL454 (Figure 6A). These trends illustrate the non-additivity (network rewiring) of parental gene expression in the progenies, where outlier trends were evident. The *OsCML27* cohorts in FL510 were enriched with salicylic acid (SA) and jasmonic acid (JA) associated genes with sustained upregulation through 144 hr (Supplemental Data Set 3). In contrast, distinct subsets of transiently upregulated genes comprised the networks in other RILs, enriched with functions associated with signal transduction, cell wall biogenesis, and growth. Transient expression among the poor RILs suggests a potential interruption of growth under severe stress. In the other poor RILs, the *OsCML27* cohorts also included downregulated genes associated with stress adaptation such as *OsFBK12* (Os03t0171600-02; Delayed senescence Kelch-repeat F-box protein), *OSISAP1* (Os09t0486500-02; Cold, drought and salt tolerance-associated A20/AN1 zinc-finger), *OsPLC4* (Os05t0127200-02; Salt and dehydration regulated Phospholipase C4), *OsNAGK1* (Os04t0550500-05; Drought-regulated N-acetylglutamate kinase-1), *OsMST6* (Os07t0559700-00; Monosaccharide transporter involved in water retention and membrane stability), and *OsHrd3* (Os03t0259300-01; Unfolded protein and apoptosis response; Supplemental Data Set 3) (Mukhopadhyay et al., 2004; Chen et al., 2013; Ohta and Takaiwa, 2015; Deng et al., 2019; Liu et al., 2019a). The superiority of the FL510 *OsCML27* network is further illustrated by the distinct upregulation of *OsIAA1* (Os01t0178500-02; auxin repressor AUX/IAA1), suggesting the interruption of auxin-mediated growth signaling in the poor RILs (Song et al., 2009).

**Figure 6.**
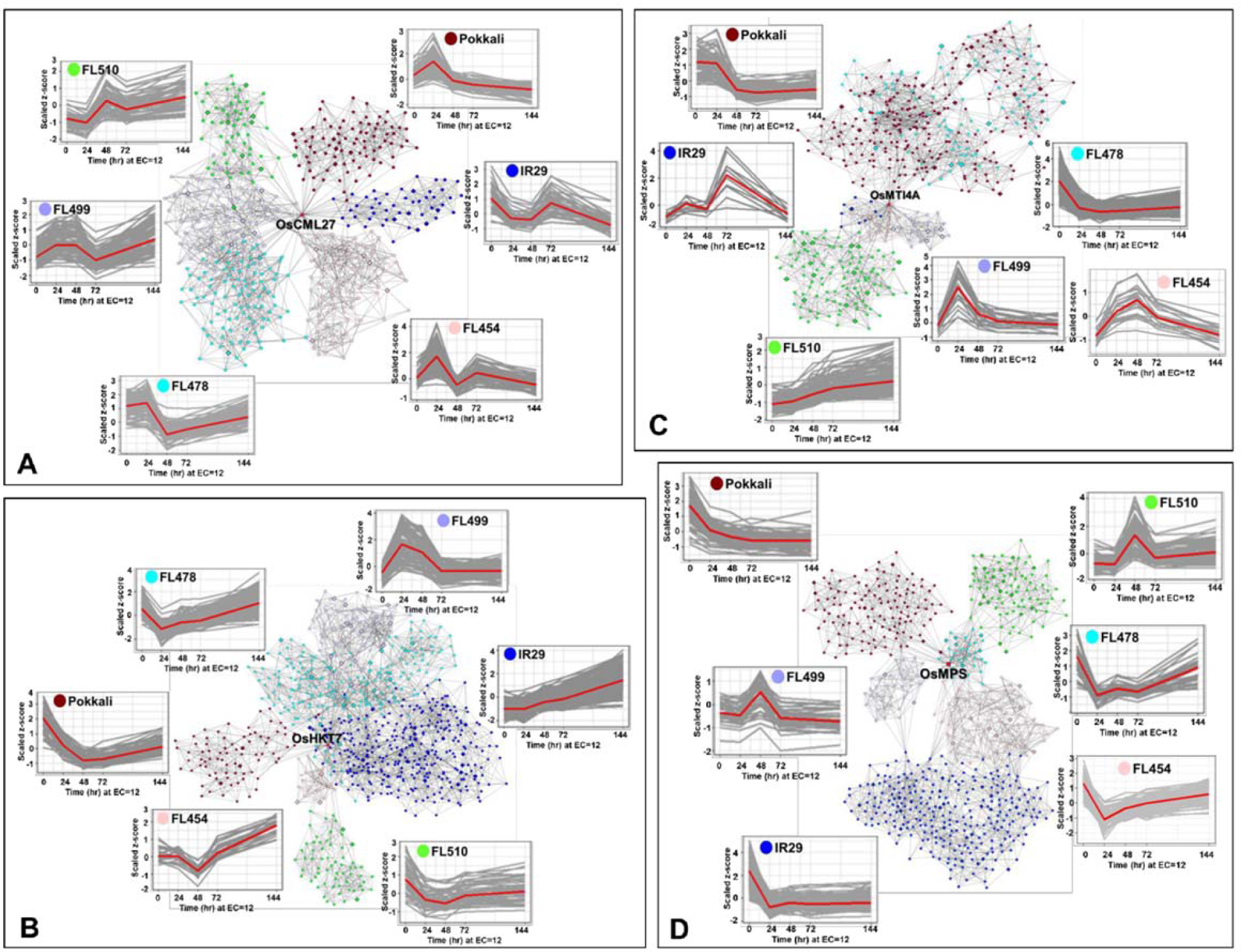
Comparison of transcriptional co-expression networks across parents and their RILs (FL454 = sensitive; FL499 = super-sensitive; FL478 = tolerant; FL510 = super-tolerant). Models represent the networks for: **(A)** *OsCML27*, **(B)** *OsHKT7*, **(C)** *OsMTI4A*, and **(D)** *OsMPS*, with important roles in developmental and stress-related responses. The temporal co-expression plots of the bait or core genes (red line) with their cohort genes (gray lines) are given for each network. Connectivity of cohort genes was based on Pearson Correlation Coefficients by mutual rank. Cohort genes are represented by nodes and coexpression is reflected on the edges.

The *OsHKT7* was upregulated in IR29 but downregulated in Pokkali (Figure 6B). The super-tolerant FL510 resembled the Pokkali profile, while the tolerant FL478 and sensitive FL454 were more similar to IR29 and the super-sensitive FL499 had non-parental profile. The *OsHKT7* cohorts had the most overlap in FL510 and Pokkali, enriched with functions associated with transcription, translation, and cell division (Supplemental Data Set 4). However, the cohorts in FL510 appeared to be uniquely enriched with cation transport functions such as *OsMTP9* (Os01t0130000-01) for manganese transport, and *OsCDT2* (Os06t0143100-01) for cadmium exclusion (Kuramata et al., 2008; Ueno et al., 2015). The cohorts in IR29, FL454, FL499 and FL478 were associated with stress sensitivity highlighted by *OsMYB30* (Os02t0624300-01) and *OsHSFB2B* (Os08t0546800-01) (Xiang et al., 2013; Lv et al., 2017).

The cysteine-rich metallothionein gene *OsMTI4A* was downregulated in Pokkali and transiently upregulated in IR29 (Figure 6C). Expression in FL478 was more similar to Pokkali, while expression in FL499 and FL454 were more similar to IR29. Most interestingly, expression in FL510 was distinct. While significant network overlaps are apparent between FL478 and Pokkali, and between IR29, FL454, and FL499, the cohort genes in FL510 were also distinct from the rest (Supplemental Data Set 5). The Pokkali and FL478 networks included downregulated genes involved in translation, cell division and growth regulation, such as CyclinF2-2 (Os02t0605000-01) and CyclinB1-5 (Os05t0493500-00) (La et al., 2006). The networks in the poor genotypes IR29, FL454, and FL499 were comprised of transiently upregulated genes involved in general stress response. The unique *OsMTI4A* network in super-tolerant FL510 was characterized by sustained co-upregulation of genes involved in hormonal, growth and stress signaling such as *BZR3* (Os06t0552300-01; repressor of brassinosteroid signaling), *OsIAA13* (Os03t0742900-01; Auxin repressor AUX/IAA13), *CDKG2* (Os04t0488000-02; cyclin-dependent kinase G-2), *OsE2F1* (Os02t0537500-01; Mitotic cycle E2F protein), and extensin (Os01t0644600-00) (Kosugi and Ohashi, 2002; Tank and Thaker, 2011; Kitomi et al., 2012; Schmitz et al., 2013; Draeger et al., 2015). The novelty of FL510 *OsMTI4A* network provides another layer of support to its unique ability to efficiently integrate stress and growth responses.

The MYB-type Multi-pass transcription factor *OsMPS* is a critical regulator of plant growth and development under stress (Schmidt et al., 2013). While this gene was significantly downregulated in both parents, different patterns of transgressive upregulation were evident across the RILs (Figure 6D). Distinct expression in super-tolerant FL510 was characterized by early induction at 48 hr. While the *OsMPS* network appeared to be distinct in each genotype, the most significant overlaps were between the super-tolerant FL510 and tolerant FL478 (Figure 6D). Cohorts in FL510 is enriched for JA and SA signaling genes such as *OsWRKY3* (Os03t0758000-01), *OsWRKY67* (Os05t0183100-01), *OsWRKY53* (Os09t0334500-01), *OsWRKY74* (Os05t0343400-01), *MYC2* (Os10t0575000-01) and *RERJ1* (Os04t0301500-01; Supplemental Data Set 6), which are important for integrating stress with development, and shared at least partially with FL478 (Ramamoorthy et al., 2008; Miyamoto et al., 2013; Ogawa et al., 2017).

The *OsMPS* network in Pokkali was enriched with genes associated with general transcription and translation, reinforcing the trends observed in its other networks. The other genotypes show few similarities with *OsCML27, OsHKT7*, and *OsMTI4A* networks with the downregulated oxidative stress genes such as *Prx19* (Os01t0787000-01; peroxidase), *Prx117* (Os08t0113000-01; peroxidase), and *OsSIK1/OsER2* (Os06t0130100-01; LRR receptor-like serine/threonine kinase), and other genes involved in cell wall biogenesis (Cosio and Dunand, 2008; Ouyang et al., 2010). These trends are consistent with the relatively high growth penalties as shown by real-time growth profiling.

Despite the similar *OsMPS* expression in the super-sensitive FL499 and super-tolerant FL510, much of the cohort genes in their respective networks were distinct. The targets of *OsMPS* that were co-upregulated in FL510 were not upregulated in FL499. This suggests a possible complete uncoupling of the upstream regulator and its target effectors in FL499, and such uncoupling may only be partial in the other genotypes. Overall, the transcriptional networks support the inherent uniqueness of FL510, due to the combined effects of parentally derived profiles and as well as those that were rewired.

The correlation coefficients across each network based on median expression values of cohort genes were used as a measure of similarity across the genotypes to further illustrate the transgressive nature of FL510 and FL499 (Figure 7A). FL510 had the lowest correlation with all other genotypes. Similarly, the transgressive nature of the super-tolerant FL510 and super-sensitive FL499 was supported by the correlation coefficients based on the number of cohort genes that overlap across the genotypes, where FL510 and FL499 tend to cluster together (Figure 7B). On the other hand, IR29 and FL478 shared many more genes amongst them than with the other genotypes. FL478 was selected for *Saltol* and for its overall similarity with IR29.

**Figure 7.**
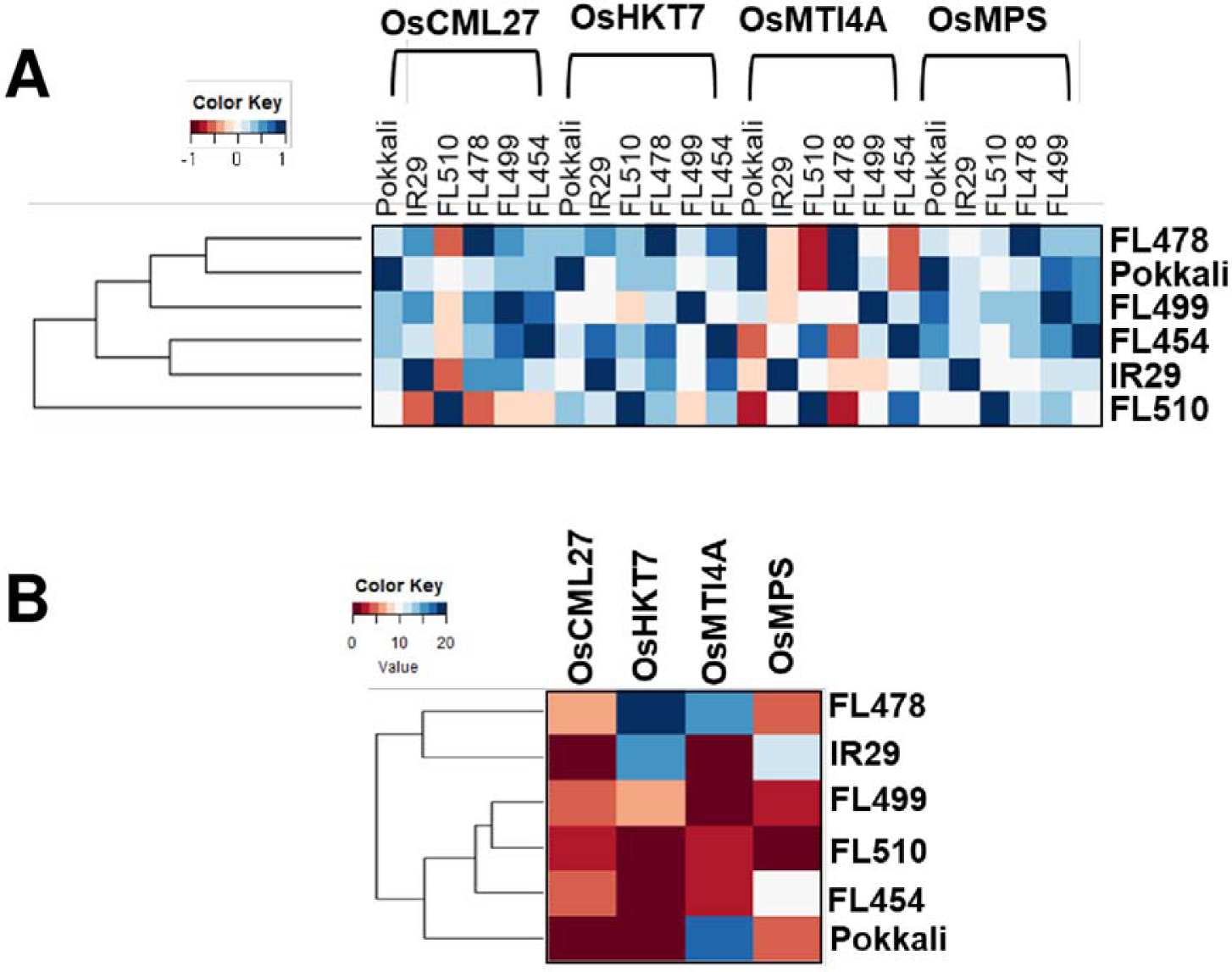
Relatedness of the genotypes according to the gene networks in Figure 6. Similarities among the genotypes were assessed by coefficient to correlation among median expression values of network components **(A)**, and by the number of shared cohort genes **(B)**.

### Transcriptome profiles as window to metabolic status

Adjustment of growth potential appeared to be critical for the superior salt tolerance of FL510. Such enhanced potential is due to the coupling of good properties from either parent or uncoupling of bad properties from the good properties from the same parent. To address this hypothesis, we assessed the similarities in primary metabolic status by reconstructing models of transcriptional status of glycolysis and tricarboxylic acid (TCA) cycles, starch metabolism, and nitrogen assimilation and transport pathways to gauge the state of plant growth. Pathway status models were based on maximum log_2_-fold change for the critical genes in each pathway.

In general, glycolysis, TCA cycle, and starch and sugar metabolism were moderately repressed in the tolerant parent Pokkali and super-tolerant FL510, in stark contrast to the inferior genotypes, especially the sensitive parent IR29 and super-sensitive FL499 where significant upregulation was quite evident (Figure 8A-C). These trends suggest that metabolic rates are faster in the poor genotypes, perhaps as consequence of the need to rapidly replenish metabolic intermediates as they are constantly depleted by cellular adjustments and defense. Superior genotypes appeared to maintain better balance and less perturbation. Nitrogen metabolic and transport pathways appeared to be enhanced in superior genotypes (Figure 8D). As nitrogen assimilation is critical for vegetative growth, these trends are consistent with the ability to maintain growth even under severe stress, as also revealed by the real-time growth profiling.

**Figure 8.**
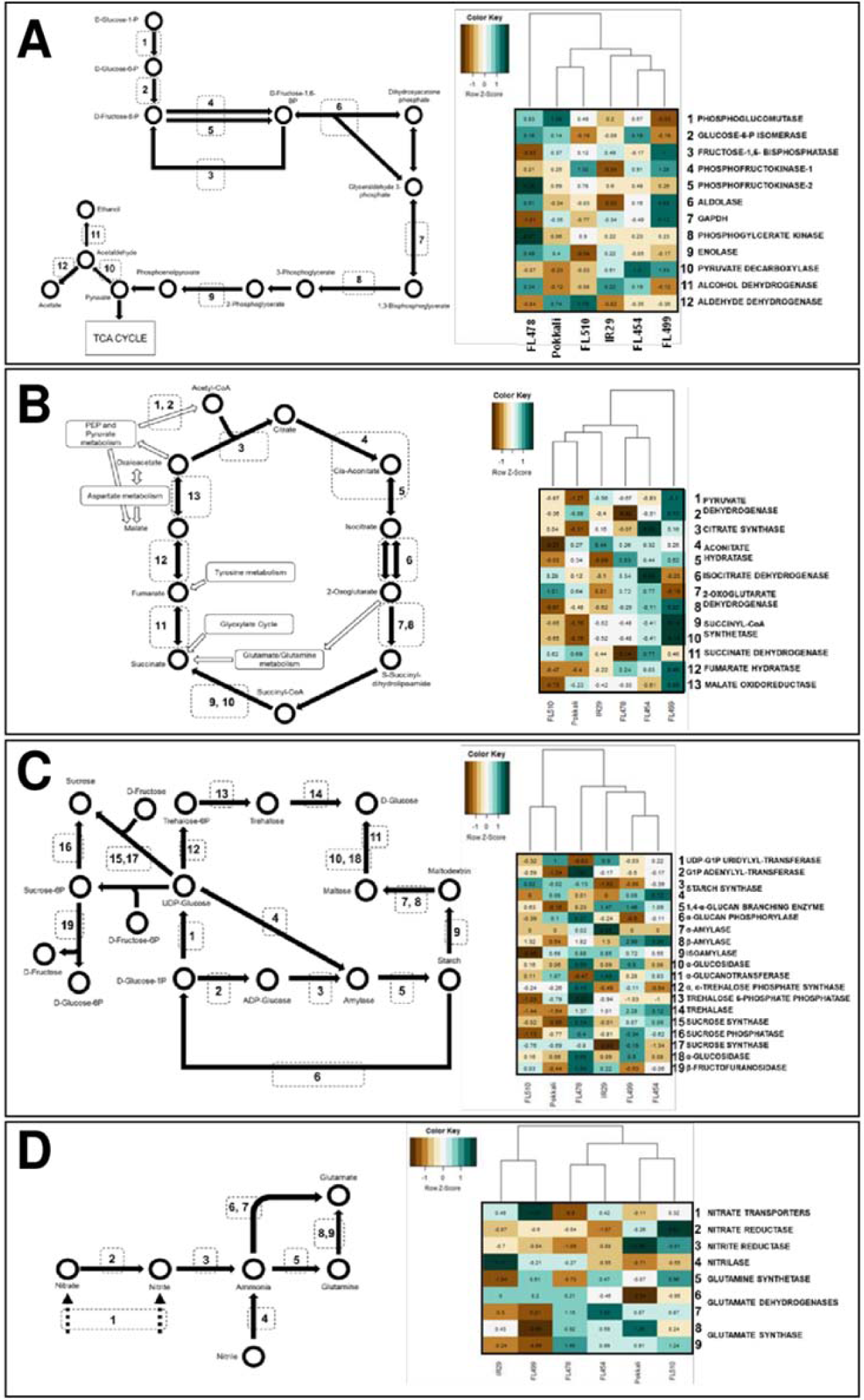
Metabolic similarities based on transcriptome profiles. KEGG models of metabolic pathways are shown on the left panel for glycolysis **(A)**, TCA cycle **(B)**, starch/sucrose metabolism **(C)**, and nitrogen assimilation **(D)**. Transcripts for enzymes in each step were mapped against each pathway. Pathway activities and the pattern of similarities across genotypes are depicted by hierarchical clustering dendrograms based on fold-change in transcript abundances at the time-point with the widest difference from control. FL510 and Pokkali share similarities in repression of the carbon metabolism pathways, while nitrogen assimilation is induced in both genotypes. In comparison, the poor genotypes cluster together.

In terms of the primary metabolic profiles, the super-tolerant FL510 was more similar to the tolerant parent Pokkali (Figure 8A-D). This trend only deviated for the Calvin cycle, where FL510 was more similar to the sensitive FL454 (Supplemental Figure 3). However, FL454 and FL510 clustered together with Pokkali, and which still points to a general slow-down or modulation of the Calvin cycle compared to other genotypes. Overall, modulation of metabolic pace in FL510 may be advantageous to maintain homeostasis under conditions when perturbed physiological state.

## Discussion

Evolutionary biology referred to the outlier phenotypes in natural populations as genetic novelties (Wagner and Lynch, 2010). Transgressive segregation is a major source of such genetic novelties, providing the raw materials for adaptive speciation under ecological niches where the majority is less adapted (Rieseberg et al., 1999). The genetic basis of transgressive segregation has been attributed to complementation and additive effects, and positive or negative epistatic interactions, among other phenomena (Dittrich-Reed and Fitzpatrick, 2013). In plant breeding, where directed mating of two individual genotypes (intra-species or inter-species) with wide phenotypic contrasts serve as foundation for stacking desired traits together, transgressive segregation could be a common occurrence. However, much of these novelties are often undetected due to small population size, narrow scope and limited resolution of phenotypic classification, limited generation time to break genetic linkage and physiological drags, and lack of sufficient data and genetic models for their prediction. Nevertheless, when interfaced with the high-resolution genetic scrutiny through genomic modeling, transgressive segregation could provide a fine-tuned combination of compatible and additive traits to address the needs for enhanced environmental resiliency of 21^st^ century crops, above and beyond what could realistically be achieved by transgenics and gene editing (de los Reyes, 2019).

In marker-assisted plant breeding, crosses between parents carrying few complementary major-effect QTL are often made to maximize genetic gains from the combining potentials of the parents, especially in the absence of linkage drags (Collard and Mackill, 2008). While much success has been achieved particularly for adaptive traits and yield potential using this paradigm, the strategy of QTL stacking is often performed with the simplistic expectations that most outcomes are positive additive effects, while ignoring much of the potential negative interactions that undermine the full force or potential of positive synergies. Negative interactions are in fact coming from the background (minor effect QTL or peripheral components) that cannot be explained by the major-effect QTL (*i.e*., Omnigenic Theory).

Focused on a well characterized genetic population of rice (IR29 x Pokkali RILs), this study was motivated by the recognition that the maximum combining potentials of parents for transgressive segregation for sustained growth under salt stress could be an equal contribution of the positive additive components (major and minor effects) from either parent, and the absence of antagonistic elements from both parents that undermine the full force of positive synergies. This study also aimed to uncover evidence of genetic network rewiring based on non-parental physiological, biochemical, and molecular attributes. The ultimate goal was to illuminate the various types of synergies that create either large positive or negative net gains.

We addressed the physiological coupling and uncoupling hypothesis by revealing that IR29 (sensitive parent) was the source of positive growth and developmental attributes that were complemented by the superior defense attributes of Pokkali (tolerant parent). Additive effects are manifested because of the absence of physiological drags that undermine the full expression of growth and developmental attributes in IR29, and the defense attributes in Pokkali either independently or through their interactions. Evidences of non-parental physiological, biochemical, and molecular attributes were uncovered in the transgressive segregants, consistent with the proposed network rewiring theory (de los Reyes, 2019).

To understand the subtleties of physiological coupling and uncoupling, we borrowed the paradigm of ‘*personalized genomics medicine*’, which considers any two individuals with similar phenotypes to have unique signatures based on multi-dimensional or holistic criteria. We scrutinize the representatives of the full phenotypic range across a recombinant inbred population at the highest resolution, which is not normally achieved in conventional plant breeding pipeline. Our results further illuminated the biological basis of transgressive segregation to complement what is known in classical plant breeding. This study provides a solid proof of concept that genetic novelties can be created by hybridization. It represents an important advance for further exploration of transgressive segregation towards novel adaptive phenotypes that may not be achieved by functional genomics, transgenics or gene editing.

### Coupling and uncoupling effects at multiple levels

The transgressive properties of the super-tolerant FL510 stems mainly from the complementation of beneficial traits (*coupling*) and shedding of potential drags (*uncoupling*) from both parents, made possible by the fortuitous recombination events and genome shuffling through at least eight generations (F_8_). Coupling and uncoupling were observed at multiple levels, *i.e*., macro-physiological, metabolome, and transcriptome. A hypothetical model is presented in Figure 9 to illustrate how the unique assemblages of properties or their opposites from each parent configured unique attributes to certain individuals.

**Figure 9.**
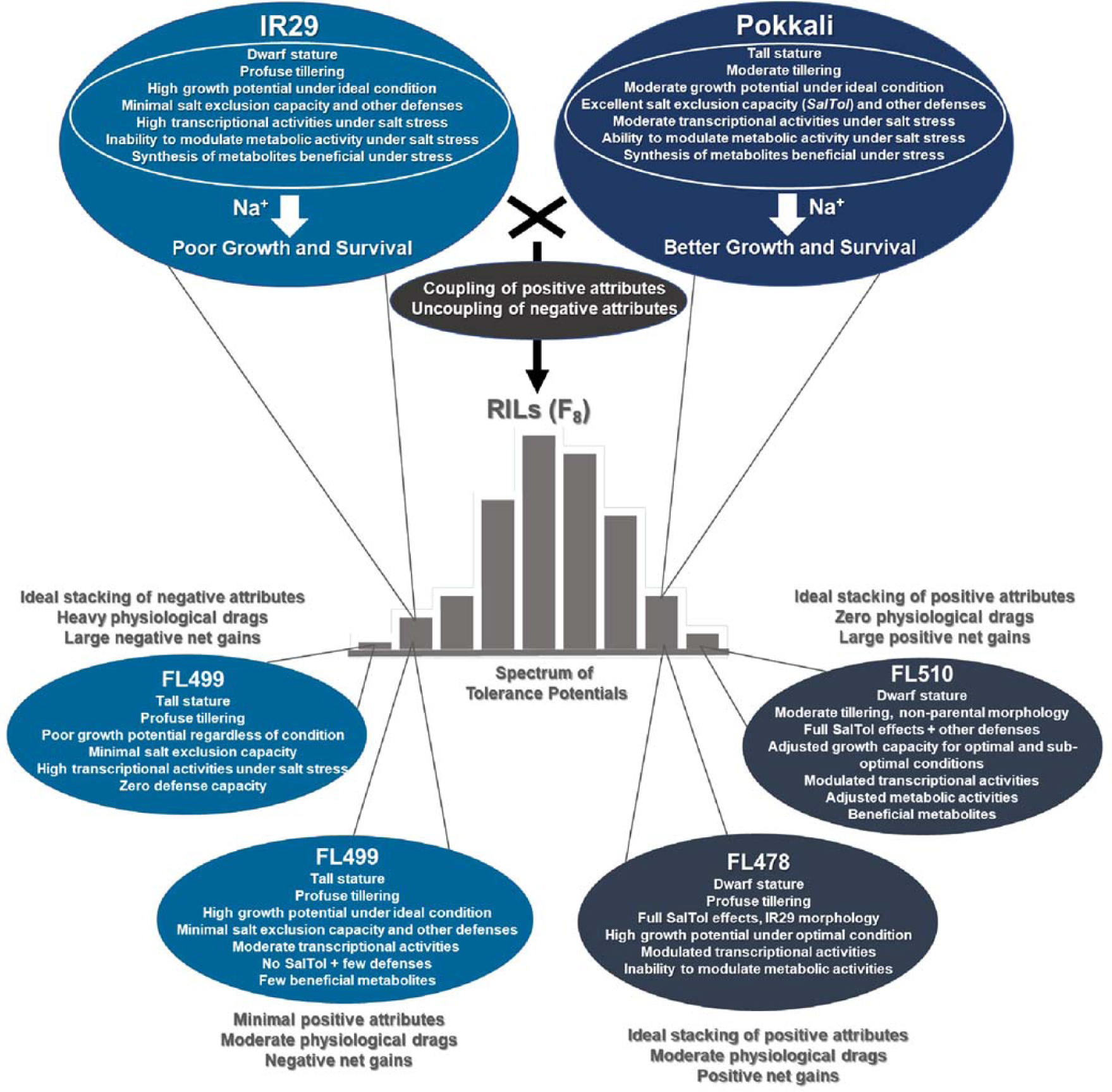
Hypothetical model of physiological coupling and uncoupling in transgressive segregants for salt tolerance across the IR29 x Pokkali recombinant inbred population based on macro-physiological, biochemical and molecular profiles according to De Ios Reyes (2019). This model proposes that the novelties of FL510 and FL499 are due to the coupling in the progeny of the good properties coming from either parent or uncoupling of bad properties from the good properties from the same parent. On top of the core mechanisms that contribute to a large proportion of phenotypic variance for defense potentials, each parent has their own characteristics that may or may not be beneficial under stress. Benefit from IR29 would be its superior growth and development potentials. Pokkali offers many stress defense mechanisms including salt exclusion. Combining the physiological potentials of parents with the reconfigured (non-parental) properties led to positive or negative coupling and uncoupling effects in RILs.

At the macro-physiological level, the best measure of variation for salt tolerance was the magnitude of growth penalty. Sub-optimal conditions slow-down the rate of photosynthesis, while requiring more resources to feed into the maintenance of homeostasis and short-term defenses (Munns and Gilliham, 2015). High-resolution phenotyping indicates similarity in the stress response capacity of FL510 and Pokkali, but a clear difference in overall performance. These genotypes had the highest aggregate phenotypic scores (APS), indicating that FL510 inherited the stress-related traits from Pokkali, most prominently the salt exclusion capacity through *Saltol* (S1A-B Figure). Evidently, the tolerant genotypes (Pokkali, FL510, FL478) clustered together when only salt exclusion is considered (Figure 2). FL478 has the *Saltol* effects yet outperformed by FL510, indicating that factors above and beyond salt exclusion maximize tolerance potential.

It is important to note that there appears to be a threshold as to how much the salt exclusion alone can effectively provide protection if other positively affecting and non-antagonistic properties are not in place. Prolonged exposure to stress will eventually damage the plant regardless of its inherent defense potential through salt exclusion. One example related to the damage incurred eventually relates to the reduction of transpiration, as water and nutrient uptake together with gas exchange still needs to happen for maintenance. Thus, salt is still inevitably transported into the plant along with its toxic effects.

Real-time growth profiling indicated that the morpho-developmental attributes inherited by FL510 from the salt-sensitive parent IR29 are positively complementing other attributes from the salt-tolerant parent Pokkali, *i.e*., synergism. For instance, smaller stature and low tillering habit distinguish FL510 and FL478 from their inferior siblings, and these traits appeared to offer fitness benefits as they are much easier to support energetically under stress when metabolic intermediates are constantly being diverted towards cellular adjustments and defenses. In contrast, the inferior genotypes FL454 and FL499 appeared to exhibit less capacity for growth modulation, as indicated by tall stature (Pokkali), profuse tillering habit (IR29), or both (FL454, FL499). Survival is prioritized over continued growth under stress; hence the growth curve plateaus prematurely. Higher PSA ratio was observed in IR29, Pokkali, and FL478 during the onset of stress compared to FL510, FL454, and FL499, possibly reflecting an attempt to increase water uptake and turgor pressure for short-term mitigation of osmotic stress.

The deviation (‘forking’) of growth curves between stress and control is a measure of how much metabolism has been hampered, and the penalty incurred due to perturbed cellular processes and injury. Being the major trade-off of survival, this deviation was minimal in tolerant genotypes especially FL510. In this context, the coupling of the morpho-developmental features of IR29 with the stress-response mechanisms of Pokkali is ideal, allowing the plant to maximize its survival responses, while minimizing the growth trade-offs by reaching the peak of vegetative growth earlier with less metabolic costs. The super-sensitive FL499 needs to cope with stress through an inferior mechanism inherited from IR29, while trying to maintain its inherent high growth capacity. This leads to a drastic decline in growth rate and extreme sensitivity to salt.

Metabolite profiles also support the idea that IR29 contributes complementary factors towards a transgressive phenotype. Metabolites were similarly expressed between FL510 and IR29, indicating beneficial factors from IR29. Meanwhile, cluster-5 indicates the ‘*drags*’ that have been uncoupled, further contributing to increased positive net gain in FL510. Cluster-4 could also represent some of the ‘*drags*’ that prevented FL478 from achieving its full potential. These results further support the hypothesis that coupled beneficial factors from both parents are essential in creating an ideal synergy and maximal net gain.

The transgressive nature of the super-tolerant FL510 is further highlighted by its unique transcriptome signatures. The robust response in FL510 represents an optimal mechanism characterized by steady and progressive upregulation or downregulation across time. Responses are either for long-term adaptation or short-term adjustment (de los Reyes et al., 2018). Gene expression in the other inferior genotypes occurred in less orderly fashion compared to the gradual but progressive nature in FL510, indicative of perturbed state rather than adaptation.

Comparison of transcriptional networks for four genes (*OsCML27, OsHKT7, OsMTI4A, OsMPS*) with important roles in development and stress also highlight the unique configuration of FL510 as the most deviant to the parental profiles, hence rewired regulatory network as an outcome of recombination. FL510 also inherited the propensity for repressed carbon metabolism and enhanced nitrogen metabolism of Pokkali. The attributes inherited from both parents led to improved potential in combination with reconfigured gene networks gained through rewiring. All these factors contributed to synergies that allow robust growth of FL510 even under salt stress, through efficient integration of developmental and stress-related responses (de los Reyes et al., 2018). In contrast, the inferior siblings have either incomplete inheritance of beneficial characters (*e.g*., FL478), stacking of detrimental characters that maintain the sensitivity properties of IR29 (*e.g*., FL454), or further stacking of physiological drags that exacerbate the baseline sensitivity inherited from IR29 (*e.g*., FL499). In the case of FL499, profuse growth was unsustainable, aggravated by the inability to adequately address the toxic effects of Na^+^.

As bottom line, the super-tolerance of FL510 and super-sensitivity of FL499 can be defined by multiple coupling and uncoupling of physiological and molecular mechanisms, hence net gains (Figure 9). In FL510, the synergism of parental attributes was evident. Most apparent is the synergy of morpho-developmental features from IR29, and salt exclusion mechanism from Pokkali. While FL478 had similar traits as FL510, it is slightly polluted with residual physiological drags that lower its overall potential, as indicated by the metabolite and transcriptome data. FL478 also had the morpho-developmental attributes of IR29 (*e.g*., tillering capacity). Evidently, sensitive genotypes are compromised by an unsustainable metabolic requirement owing to their very active growth, while lacking the effective stress response mechanisms of Pokkali.

At the transcriptome level, the ideal synergism in FL510 allows for a sustained response through the stress period. In contrast, inferior genotypes must respond immediately, leading to an overreaction, which is energetically wasteful and detrimental in the long-term. If the stress is not relieved, the net result is a deficit that may not be reconciled, thus large penalties to survival and growth. The chances that extreme phenotypes such as those in FL510 and FL499 would occur in a population are small, as there needs to be a specific combination of parental traits and creation of novel regulatory networks to create an ideal or non-ideal synergy.

### Relevance of the coupling and uncoupling effects to the Omnigenic Theory

The Omnigenic Theory represents a modern paradigm for explaining the genetic causes of complex diseases in humans that cannot be fully explained by polygenic effects (Boyle et al., 2017). According to this theory, complex traits are controlled by thousands of genes or loci scattered throughout the genome, rather than a small group of genes detected by conventional QTL mapping or genome-wide association. Core genes/loci are central to the trait and detected as QTL or major genes by forward genetic screens. The peripheral genes/loci are below the threshold of QTL detection, as they have immeasurable effects as individual components of a network. However, their combined effects can be equal to or more than the effects of the cores (Liu et al., 2019b). To maximize the potential for trait expression through G x E, both the combinations of the core and peripheral alleles and their interactions must be optimal.

Transgressive segregants are consequences of the complementation of compatible traits from both parents (Rieseberg et al., 1999). Their novelties are also due to the absence of physiological or genetic linkage drags. However, if fortuitous, uncoupling events could also result in the loss of positive synergies (de los Reyes, 2019). This study illustrated that complementation in either synergistic or antagonistic fashion creates transgressive properties at both sides of the phenotypic spectrum, extending beyond the typical salt exclusion mechanism contributed by Pokkali. In FL510, growth reduction was minimized since metabolic impairment was not as extensive compared to other genotypes. FL478 and Pokkali both possess salt exclusion mechanisms; however, they lack the multitude of complementary but minute effects for further incremental enhancements.

In FL510, the synergism among the positive parental attributes buoy plant health enough to survive under stress. Salt exclusion mechanism reduces Na^+^ toxicity in metabolically active tissues, which works together with the lower metabolic requirement for growth and development, hence a robust response. Transport of Na^+^ and Cl^-^ alone constitutes major cellular energy costs. When minimized, it allows the plant to survive longer under saline environment (Tyerman et al., 2019; Munns et al., 2020). The collective responses uncovered in FL510 are consistent with the Omnigenic Theory, as no single mechanism is adequate to explain its extreme phenotype. The same thing is true for the super-sensitive FL499 with antagonistic effects. Physiological gain is the result of ideal synergy, reflecting the concept that the majority of the genome contributes to the full physiological potential of each individual in a population.

### Comprehensive phenotyping and genomic modeling of transgressive traits

The transgressive salt tolerance of FL510 may not extend to other types of stress such as drought. Indeed, other RILs derived from IR29 x Pokkali have been shown to survive under severe water-deficit better than FL510 (Gendron, 2019). While cases of transgressive properties such as those observed in FL510 and FL499 may be rare in a population, individuals that exceed the parental range may be more common than perceived. Repeated recombination events can create incremental improvements that allow individuals to have slight advantages over their parents. Often, individuals that display such properties are cast aside, as they do not offer much advantage in a constrained plant breeding pipeline. These individuals also often carry undesirable ‘linkage drags’. However, these individuals offer invaluable tools in modeling how transgressive phenotypes arise in a population.

Genomic selection promises to identify desirable but rare recombinants for traits governed by complex multi-loci interactions by taking into consideration the core and peripheral components and all forms of interactions (de los Reyes, 2019). Increasing the accuracy of modeling is dependent on the number of individuals in the modeling population. With enough individual data points, a wider coverage of possible permutations of recombinants can be mined for genomic patterns that lead to transgressive phenotypes. Thus, the interaction networks from both core and peripheral components may be modeled for their individual weights.

It must be remembered that the process from which transgressive individuals arise in plant breeding is similar in many ways to the process by which new phylogenetic lineages could develop in natural populations through hybridization (Rieseberg et al., 1999; Dittrich-Reed and Fitzpatrick, 2013). The fact that transgressive individuals can arise through repeated recombination events is a hopeful observation. All progeny in a cross is potentially useful for modeling transgressive phenotypes. Each recombination event adds a possible genomic permutation, and these should not be discarded, as various types of synergism can be identified and understood from such individuals.

It is also important to increase the resolution of analysis with multiple layers of data across a population. A multi-tier view of phenotypic range will allow a more comprehensive integration of the ‘*omics*’ data to unravel the wirings of transgressive phenotypes, analogous to the paradigm of *personalized genomic medicine*. This study represents a serious attempt to test this approach that is unconventional even in modern plant breeding pipelines, *i.e*., analyzing multiple layers of data to decipher complementary or antagonistic effects. While it has limitations, future advances in technology will allow a more seamless data integration. Addition of other layers, such as epigenomic configuration and chromatin structure will certainly enhance the prospect of modeling (de los Reyes, 2019). Additionally, beneficial traits do not come solely from one parent, and the value of a genotype as donor cannot be predicted based on phenotype alone, unless its combining potential has been examined in recombinants. Extensive phenotyping may reveal cryptic characters that are beneficial but are often overshadowed by other negative traits.

## Materials and Methods

### Phenotypic evaluation of the RIL population

The RILs of IR29 (*Xian/indica*; salt-sensitive) x Pokkali (*Aus*; salt-tolerant) consisted of 123 individuals as the core QTL mapping population. Segregation for salt tolerance was established earlier at IRRI based on Standard Evaluation Score (SES) and shoot Na^+^/K^+^ at V1 to V5 for the fine-mapping of *Saltol* at EC= 9 dS/m (Singh et al., 2007; Thomson et al., 2010). Based on IRRI’s data, a comparative panel of 67 individuals representing the full range of SES and Na^+^/K^+^ was subjected to a multi-parameter evaluation at V4 to V12. The replicated (n=8) experiments were conducted in hydroponics at 30-35°C day, 24-26°C night; 20% to 30% RH; 12-hr photoperiod with 500 μmol m^−2^s^−1^ light intensity. Physiological parameters included SES (Gregorio et al., 1997), Na^+^/K^+^, electrolyte leakage index (ELI) (Ballou et al., 2007; de los Reyes et al., 2013), peroxidase (POX) activity, lipid peroxidation (LP) (Jambunathan, 2010), and shoot biomass.

Seedlings were grown in standard peat moss for fourteen (14) days. Individual plants were transplanted to 0.6-gallon hydroponic buckets with 1 g/L Peter’s Professional 20-20-20 General Fertilizer (JR Peters Inc., Allentown, PA) at pH 5.8, supplemented with 0.4 g/L FeSO_4_·7H_2_O. Plants were subjected to salt stress at tillering stage (V4 to V12; (Counce et al., 2000) with ∼120mM NaCl (EC= 12 dS/m). Samples for physiological assays, RNA extraction, and biomass measurements were collected at 0 hr (control) and 24, 48, 72, and 144 hr after stress.

### Shoot biomass ratio, stomatal conductance, and standard evaluation score (SES)

Shoot fresh weight was measured from three random plants for each genotype at 0 hr and 144 hr after stress. Tissue samples were oven-dried (50°C) for five days and weighed to calculate the % biomass. Biomass ratio >1 indicates growth, while ratio <1 indicates penalty. Stomatal conductance was measured using the SC-1 Leaf Porometer (Meter Group Inc., Pullman, WA). These measurements were taken at 0 hr and 144 hr and presented as control/stress ratio. Stomatal conductance was measured at the same time of the day to prevent diurnal effects. At the end of the stress treatments (17 days), the overall health of each plant was rated visually using IRRI’s Standard Evaluation System (SES) (Gregorio et al., 1997). SES is based on a ranking of 1 to 9 with higher score indicating more severe magnitude of stress injury.

### Electrolyte leakage index

Leaf Electrolyte Leakage Index (ELI) was used as a function of cellular injury from osmotic stress as well as ion uptake according to Ballou et al. (2007) and de los Reyes et al. (2013). Briefly, leaf discs were sampled from individual plant replicates (n = 5) and placed in 5 mL ultrapure water (18 megaOhms). Electrical conductivity (EC) was measured with a conductivity meter (Thermo Scientific, Waltham, MA) after allowing the tissue electrolyte to leach out (stress EL). Total tissue electrolytes were measured after boiling at 95°C. Electrolyte leakage was calculated as: ***EL***_***sample***_***= EC***_***boiled***_***/EC***_***unboiled***_ ***x 100***. ELI was calculated with the following equation and expressed as a mean of ELI (n = 5): ***ELI = EL***_***144hr***_***/EL***_***0hr***_ ***x 100***.

### Peroxidase activity assay

Total peroxidase activity (POX) was measured with the Amplex® Red Hydrogen Peroxide/Peroxidase Assay Kit according to manufacturer’s instructions (Invitrogen, Carlsbad, CA). Pulverized leaves were homogenized in 20mM sodium phosphate buffer (pH 6.5) and centrifuged at 9000 x g for 10 min at 4°C. Samples were reacted with the Amplex® Red mix in the dark for 30 min, and absorbance at 560 nm was presented as means (n = 3). POX was based on a standard curve of pure horseradish peroxidase at a range of 0 to 2 mU/mL.

### Lipid peroxidation assay

Assays for lipid peroxidation were adopted from Jambunathan (2010). Leaf samples were pulverized in liquid nitrogen, homogenized with 0.1% trichloroacetic acid (TCA), and centrifuged at 10,000 x g for 15 min at room temperature. The supernatant was combined with 20% TCA and 0.5% thiobarbituric acid (TBA) and incubated at 95°C for 30 min. Absorbance was measured at 530 nm and 600 nm, with A_600_ serving as a background absorbance against A_532_. Values were used to determine the malondialdehyde (MDA) content as an estimate of broken lipid membranes reacting with TBA. MDA content was based on Beer-Lambert equation: ***C* = *A*/ *ε x l***, where *A* = differences in absorbance at 530 nm and 600 nm, ***ε*** = extinction coefficient of 155 mM^-1^cm^-1^, *l* = length (cm) of light path, and *C* = content (mM) of MDA.

### Na^+^ and K^+^ quantification

Na^+^ and K^+^ contents of pulverized tissues were determined by nitric-perchloric acid digestion (AOAC, 1990), measured on an AA unit per Western States Version 4.00 P-4-20 (A&L Plains Analytical Laboratory, Lubbock, TX). Values were presented as % Na^+^ or K^+^ per gram of sample (n = 3). Total Na^+^ and K^+^ were also determined for image-based phenotyping experiments. Pulverized tissues were digested with dilute HNO_3_ (0.5M) at room temperature and the supernatant was used for flame photometry using the Model 420 Flame Photometer (Sherwood Scientific Ltd., Cambridge). Na^+^ and K^+^ concentrations were calculated based on standard solution with 0.5/1 mM NaCl:KCl using the same sample dilution factor, amount of digestion solution, and initial sample weight as follows: [Na^+^]/[K^+^] = Reading/100 x Standard/100 x Digestion solution x Dilution factor/Dry weight x 100 (n = 5).

### Proline content

Total proline content of leaves (V7 to V9) was determined according to Bates et al. (1973). Briefly, pulverized leaf tissues (100 mg) were homogenized in 5 ml sulfosalicylic acid (3% w/v) and then centrifuged at 13,000 x g for 10 min to collect the supernatant. The assay solution was comprised of fresh ninhydrin (2.5% w/v), glacial acetic acid (60% v/v), and phosphoric acid (40% v/v). Supernatant (0.1 ml) was combined with 0.2 ml assay solution and 0.2 ml glacial acetic acid and was incubated at 95°C for an hour. Chromophore was extracted with 1 mL toluene, and the organic phase (0.1 ml) was recovered into 96-well microtiter plates for absorbance measurements at 520 nm. Quantification was based on a standard curve of 0 to 100μM pure proline (Fisher BioReagents, BP392-100). Assays were performed with replicates (n = 3).

### Aggregate phenotypic scores (APS)

Values from each parameter were transformed into relative values on a scale of 1 to 10 across the RIL population. The highest value was set to a score = 10, while the lowest value was set at score = 1 according: ***y = 1 + (x – A) x (10 – 1)/ B – A***, where y = normalized value, x = raw value, A = minimum, and B = maximum. Scores in each genotype were combined into a single normalized phenotypic score and compared to SES. Scoring matrix across the RILs was analyzed by hierarchical clustering using the ‘pvclust’ package in R (Suzuki and Shimodaira, 2006).

### Real-time plant growth profiling

The comparative panel comprised of IR29 (sensitive parent), Pokkali (tolerant parent), FL478 (tolerant RIL), FL510 (super-tolerant RIL; positive transgressive), FL454 (sensitive RIL), and FL499 (super-sensitive RIL) were subjected to digital growth profiling under control and stress with the LemnaTec Scanalyzer 3D platform at the University of Nebraska, Lincoln, NE. Disinfected seeds with were germinated in 0.5X Murashige-Skoog (MS) agar for five days. Seedlings were sown in hydroponics consisting of Turface MVP® submerged in Yoshida solution (Yoshida et al., 1971). Plants were loaded onto the LemnaTec Scanalyzer 3D platform after 14-days. Salinity treatment was introduced by adding NaCl (270 mM NaCl:9.9 mM CaCl_2_) to EC= 4.5 dS/m (45mM). This initial salt treatment was escalated the next day to EC= 9 dS/m (90mM) and maintained at that level for the duration of the experiment.

Solutions were changed every other day to maintain nutrient content and salinity. Plants were digitally imaged daily for eighteen (18) days with an RGB and hyperspectral camera. The RGB images were analyzed through a Matlab application, and PhenoImage, (University of Nebraska-Lincoln). Plant size was initially reported as the number of pixel squares, which were then converted into cm^2^ using the Fiji software (Schindelin et al., 2012). Plant height was reported as measured by PhenoImage. Hyperspectral image variances were computed using Matlab.

### Metabolite profiling by LC-MS/MS

Shotgun metabolite analysis was performed by liquid chromatography with tandem mass spectrometry (LC-MS/MS) at the Texas Tech University Center for Biotechnology and Genomics. Pulverized leaf samples (100 mg) was homogenized with chilled chloroform:methanol:water (1:2.5:1 [v/v/v]) and used for the separation of aqueous and lipid phases. Samples were resuspended in 0.1% formic acid (v/v) and 5 μl aliquots were used for LC-ESI-MS/MS using the Dionex Ultimate 3000 nano-LC (Thermo Scientific, San Jose, CA) interfaced to Q Exactive™ HF Hybrid Quadrupole-Orbitrap™ mass spectrometer (Thermo Scientific, San Jose, CA). Metabolites were separated on a C18 Acclaim PepMap RSLC column (Thermo Scientific, San Jose, CA) at constant flow of 0.3 µL/min with mobile phase solvent-A (97.9% water/2% ACN/0.1% FA) and solvent-B (99.9% ACN/0.1% FA). Metabolites were separated by following the gradient of solvent-B. For MS/MS analysis, the first scan was 50–500 m/z at mass resolution of 120,000. The ten most intense ions from the first scan were used for HCD MS/MS with elevated collision energy of 20-60% in positive and negative modes.

The Compound Discoverer v3.0 (Thermo Scientific) was used to detect compounds with “Predicted Formula”, followed by automatic online library search against the mzCloud and ChemSpider databases (Pence and Williams, 2010; Mistrik, 2018). Compounds from mzCloud library was checked using the mirror plot of MS/MS spectra in library standards. For quantitative analysis, area under peak was calculated with: precursor ion mass tolerance = 5 ppm, intensity tolerance = 30%, minimum peak intensity = 1×10^6^, alignment mass tolerance = 5 ppm, peak alignment maximum shift = 2 min, and ass tolerance of fragment ion= 10 ppm.

### RNA-Seq and transcriptional network modeling

Total RNA was extracted from frozen leaf tissues using miRVana™ miRNA Isolation Kit (Invitrogen, Carlsbad, CA) and used to construct time-course (0, 24, 48, 72, 144 hr) RNA-Seq libraries. Strand-specific 150-bp paired-end RNA-Seq libraries were sequenced on Illumina HiSeq3000 with two replicates (Oklahoma Medical Research Foundation, OK). RNA-Seq data were analyzed using the established pipeline (Kitazumi et al., 2018). Sequence output from indexed RNA-Seq libraries were preprocessed with Cutadapt (Martin, 2011) and mapped against the Kasalath reference (Sakai et al., 2013) using HISAT2 (Kim et al., 2015). Transcript read counts were normalized by TMM (Trimmed Mean M-values) and differential expression was examined by edgeR using a false detection rate of 0.05 (McCarthy et al., 2012).

Gene expression clusters were identified and visualized using the ‘MBCluster.Seq’ package in R (Si et al., 2013). Transcript abundance were investigated by the mutual rank (MR) method (Obayashi and Kinoshita, 2009). Pearson’s correlation coefficient (PCC) matrix was established for each cluster, and genes with PCC < 0.95 were excluded. MR was calculated through: ***MR***_***AB***_ ***= Square-root of Rank***_***A to B***_ ***x Rank***_***B to A***_. Gene interactions with values < 10 were excluded. Genetic networks were constructed and visualized using Cytoscape (Smoot et al., 2010). The annotation applied to RNA-Seq dataset was based on UniProt with additional expansion from other relevant databases (The UniProt Consortium, 2018). Metabolic pathway status modeling was aided by the Kyoto Encyclopedia of Genes and Genomes (KEGG) (Kanehisa and Goto, 2000; Kanehisa et al., 2018).

### Statistical analysis

Statistical analyses were conducted with R 3.5.2 (R Core Team, 2020). Pearson’s correlation coefficient matrices, gene ranks, and mutual ranks for genetic network and PCA were scripted in R-package. Tukey’s HSD post-hoc tests following ANOVA were performed with the ‘agricolae’ package (de Mendiburu and de Mendiburu, 2019). K-means clustering was through the K-means++ code included in ‘mytools’ package (Yasumoto, 2020). Data visualization were through the ‘ggplot2’ package (Wickham and Chang, 2008).

### Accession Numbers

The RNA-Seq data used in this manuscript are publicly available in the NCBI Short Read Archive (PRJNA378253:SRR11528269-SRR11528295).

## Abbreviations

QTL: quantitative trait loci;
RIL: recombinant inbred line;
SES: standard evaluation score;
ELI: electrolyte leakage index;
LP: lipid peroxidation;
POX: total peroxidase activity;
APS: aggregate phenotypic score;
NPT: new plant-type;
PCA: principal component analysis;
SA: salicylic acid;
JA: jasmonic acid;
TCA cycle: tricarboxylic acid cycle

## Acknowledgements

This work was supported by NSF-IOS Plant Genome Research Program Grant-1602494 and Bayer CropScience Endowed Professorship. Genomic computations were performed using the supercomputing facilities at the ROIS National Institute of Genetics, Mishima, Japan, and Texas Tech University High-Performance Computing Cluster. Metabolomic experiments were performed at the Texas Tech University Center for Biotechnology and Genomics. Digital phenotyping was performed using the LemnaTec facilities at the University of Nebraska-Lincoln. We thank Dr. Gurdev S. Khush for his valuable suggestions on the manuscript. We dedicate this work to the late Dr. Darshan S. Brar, whose far-reaching vision and ingenuity served as inspiration to this project.

## Author Contributions

ICMP and BGDR designed the experiments and wrote the manuscript. ICMP, AK, KRC, BP, MZM, and HW performed the experiments and analyzed the data. RKS and GBR generated the RIL population. BGDR conceptualized the whole project.

---

**Supplemental Figure 1.**
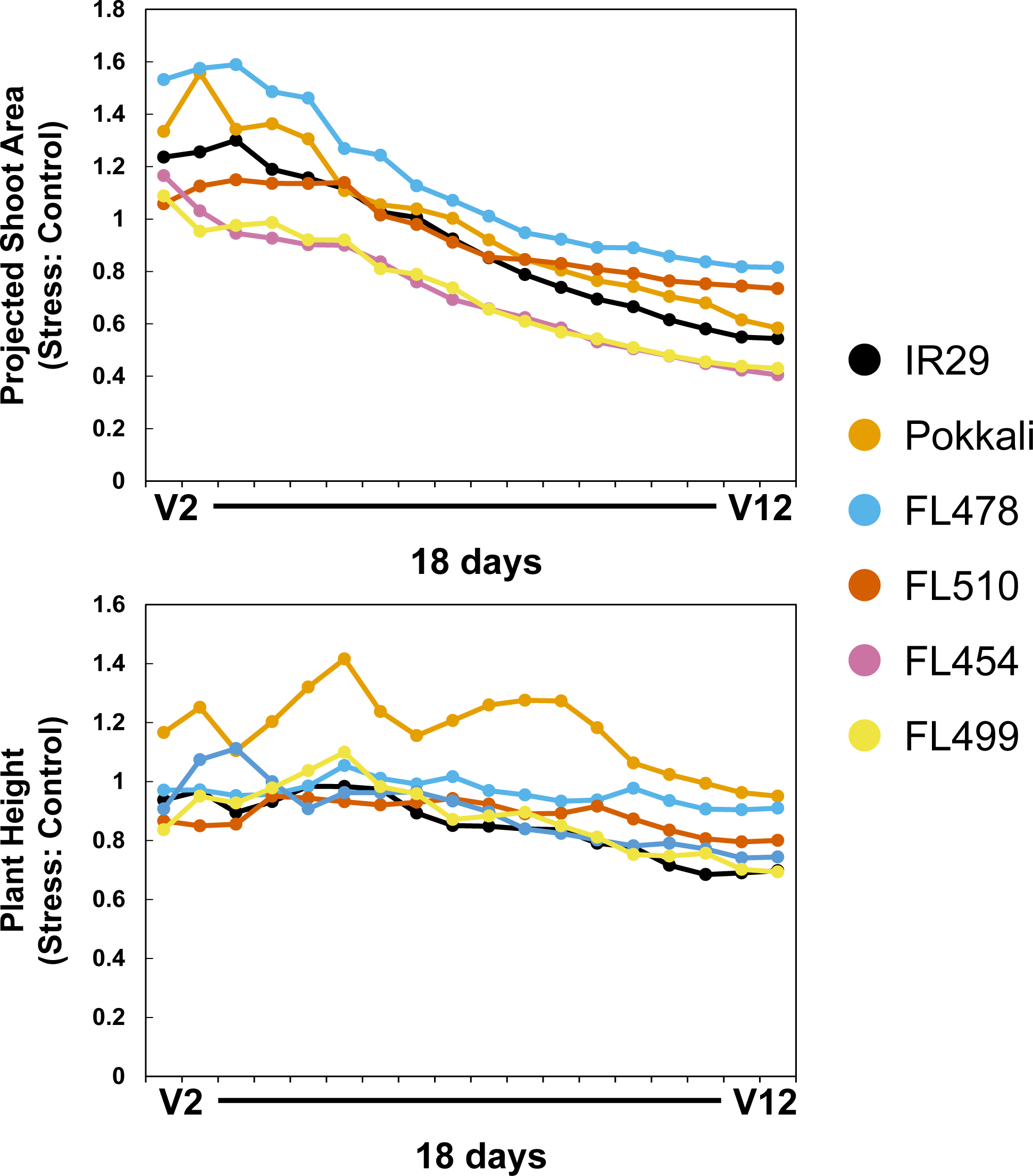
Stress to control ratios of projected shoot area (PSA) and plant height among the different genotypes used in the study across the real-time imaging period. Ratios of stress and control measurements of PSA and plant height were plotted for all the genotypes used in the study in the same time frame as Figure 1. This was used to assess the extent of growth penalty incurred by each genotype through the stress period. Ratio values greater than 1 indicate a larger value for the stress treatment compared to the control, while those lower than 1 indicate a reduction in growth relative to the control.

**Supplemental Figure 2.**
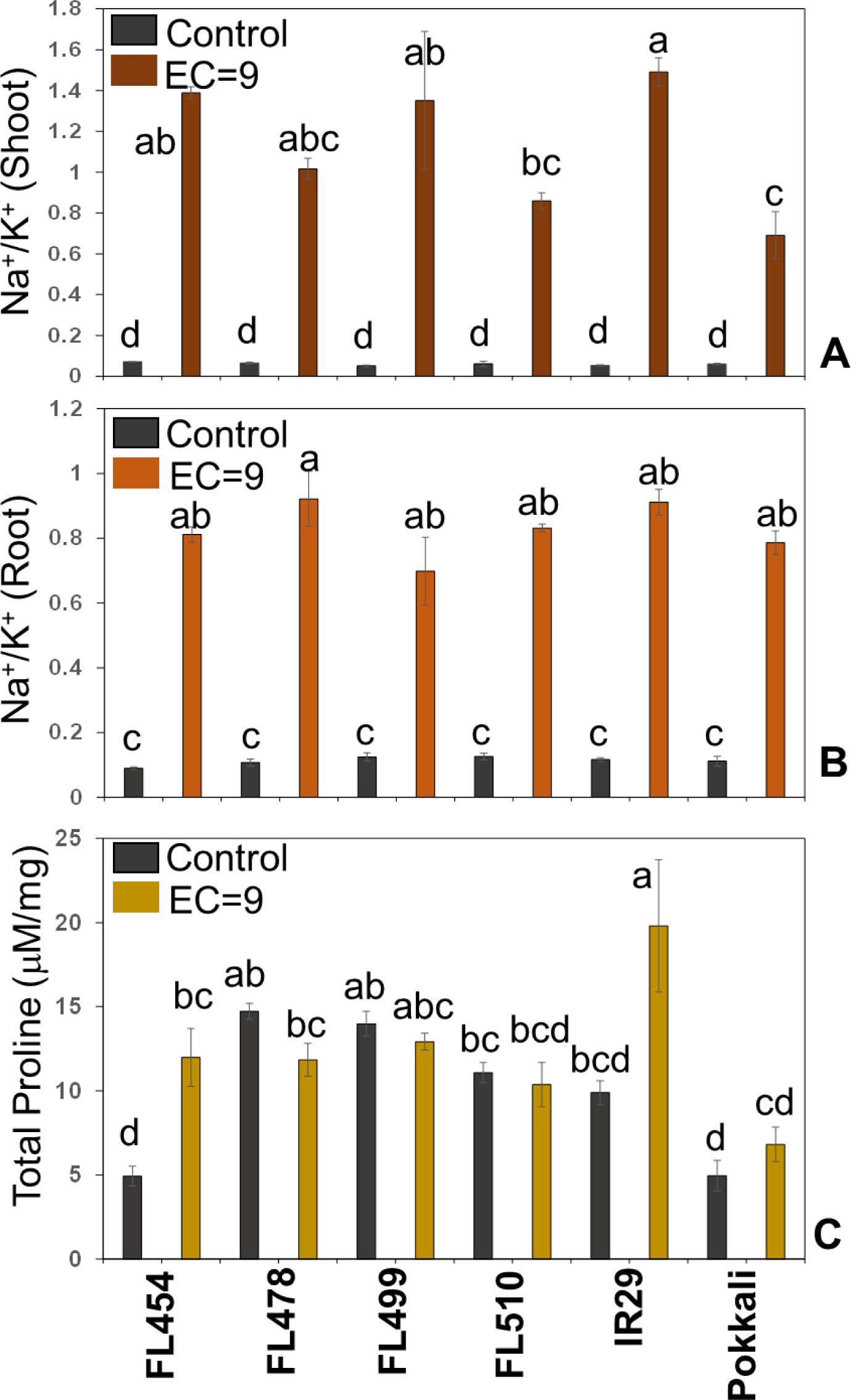
Physiological characterization of the representative genotypes used in the real-time growth profiling experiment at EC= 9. Physiological indicators of cellular defense and adjustment potentials during the 18-day period at EC=9 were evaluated including Na^+^/K^+^ in the shoot (A), Na^+^/K^+^ in the root (B), and total proline content in leaves (C). For the Na^+^/K^+^ analysis (A and B), bar graphs represent the mean Na^+^/K^+^ (n = 5) with standard errors. One-way ANOVA with an HSD test (α = 0.05) was used to determine significant differences between genotypes and treatments. Lower-case letters signify treatment groups, separated by significant mean differences, with ‘*a’* representing the group with the highest means. For the proline content analysis (C), bar graphs represent the means of total proline in leaves (n= 3) with standard error bars. One-way ANOVA with HSD test (α = 0.05) was used to determine significant differences between the different treatments. Lower-case letters signify treatment groups separated by significant mean differences, with ‘*a’* representing the group with the highest means.

**Supplemental Figure 3.**
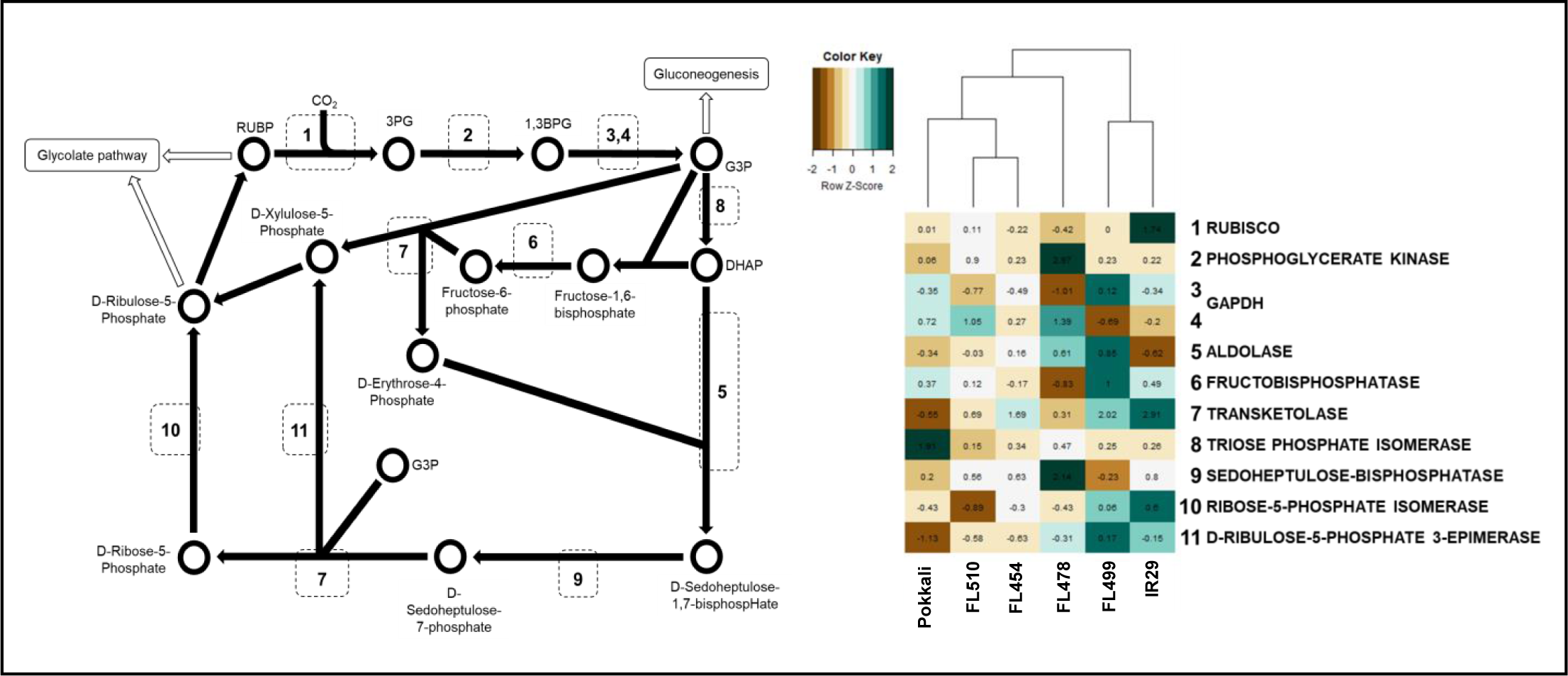
Pathway induction heatmaps under stress for genes in the Calvin cycle. Values were generated from the fold-change under control conditions and the stress time point with the widest margin from control. The genotypes were hierarchically clustered to show similarity in pathway induction. RuBP = Ribulose 1,5-bisphosphate; RUBISCO = Ribulose-1,5-bisphosphate carboxylase/oxygenase; 1,3-BPG = 1,3-bisphosphoglycerate; PPK = Phosphoglycerate kinase; 3-PG = 3-phosphoglycerate; GAPDH = Glyceraldehyde-3-phosphate dehydrogenase; GAP = Glyceraldehyde-3-phosphate; TPI = Triose-phosphate isomerase; DHAP = Dihydroxyacetone phosphate; G3P = Glyceraldehyde 3-phosphate.

**Supplemental Table 1.**
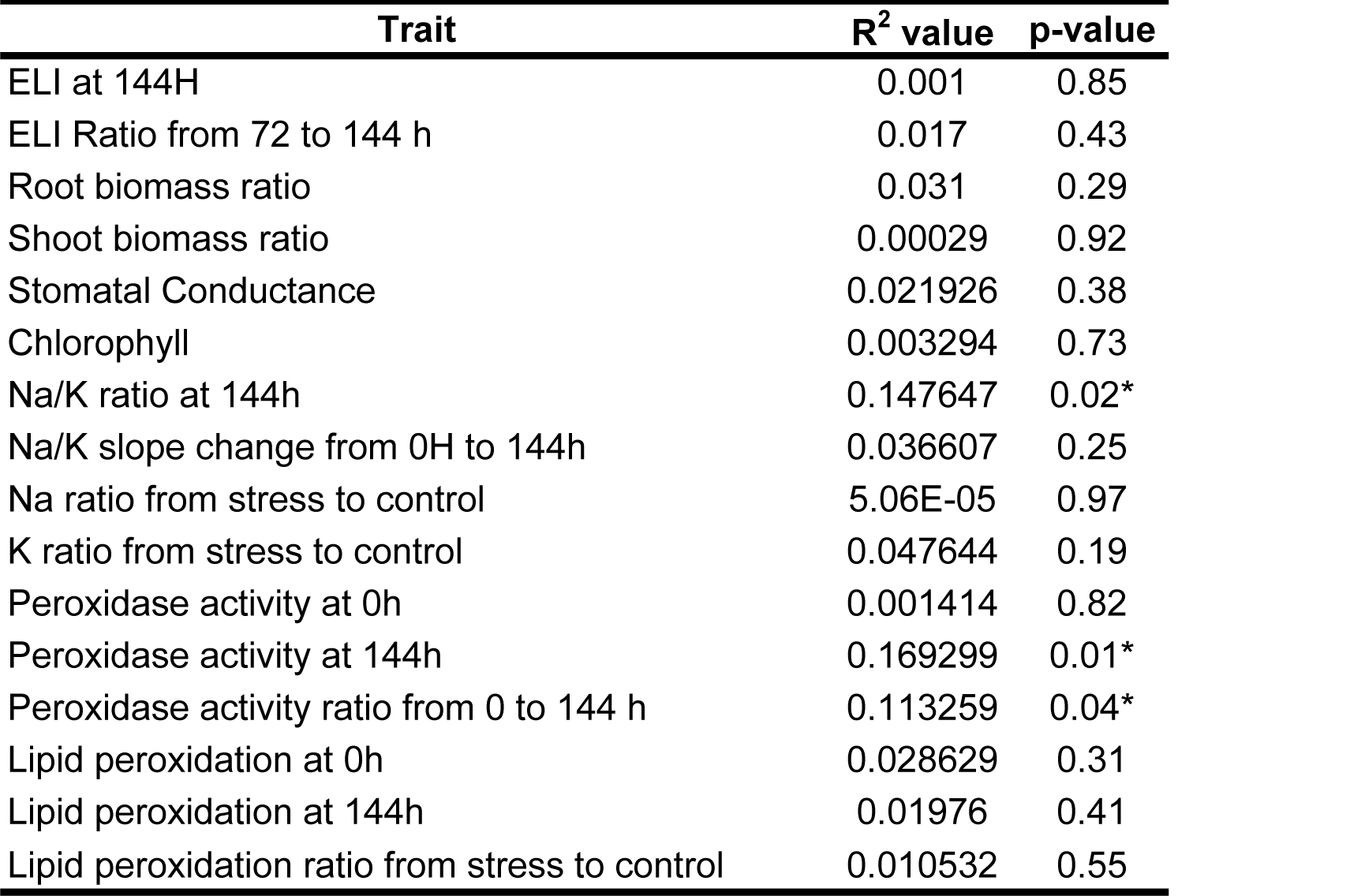
R^2^ and P-values of different traits tested in the phenotyping of the IR29 x Pokkali RIL population relative to SES. Traits with significant relationship with SES based on regression analysis (P < 0.05) are marked with an asterisk (*).

| Trait | R<sup>2</sup> value | p-value |
| --- | --- | --- |
| ELI at 144H | 0.001 | 0.85 |
| ELI Ratio from 72 to 144 h | 0.017 | 0.43 |
| Root biomass ratio | 0.031 | 0.29 |
| Shoot biomass ratio | 0.00029 | 0.92 |
| Stomatal Conductance | 0.021926 | 0.38 |
| Chlorophyll | 0.003294 | 0.73 |
| Na/K ratio at 144h | 0.147647 | 0.02* |
| Na/K slope change from 0H to 144h | 0.036607 | 0.25 |
| Na ratio from stress to control | 5.06E-05 | 0.97 |
| K ratio from stress to control | 0.047644 | 0.19 |
| Peroxidase activity at 0h | 0.001414 | 0.82 |
| Peroxidase activity at 144h | 0.169299 | 0.01* |
| Peroxidase activity ratio from 0 to 144 h | 0.113259 | 0.04* |
| Lipid peroxidation at 0h | 0.028629 | 0.31 |
| Lipid peroxidation at 144h | 0.01976 | 0.41 |
| Lipid peroxidation ratio from stress to control | 0.010532 | 0.55 |

**Supplemental Table 2.** Summary and mapping statistics of the RNA-seq libraries constructed for the parental genotypes and selected RILs.

| Sample | Replicate | Number of Paired Reads |  |  |  |  |  |
| --- | --- | --- | --- | --- | --- | --- | --- |
|  |  | IR29 | Pokkali | FL478 | FL510 | FL499 | FL454 |
| 0h | A | 16,736,902 | 21,972,858 | 19,519,378 | 27,741,440 | 25,303,994 | 22,958,328 |
| 0h | B | 23,566,931 | 31,223,873 | 27,374,472 | 31,095,774 | 28,887,331 | 32,035,095 |
| 24h | A | 17,387,989 | 18,977,029 | 16,264,496 | 30,592,532 | 25,009,427 | 17,272,958 |
| 24h | B | 24,939,854 | 27,237,708 | 23,010,419 | 34,146,243 | 28,503,630 | 24,233,186 |
| 48h | A | 31,555,196 | 26,903,719 | 28,145,394 | 31,562,536 | 27,070,597 | 31,891,918 |
| 48h | B | 33,789,866 | 30,030,083 | 30,990,592 | 34,024,720 | 30,210,218 | 34,196,118 |
| 72h | A | 25,291,270 | 30,502,192 | 27,587,928 | 26,650,239 | 25,432,219 | 30,055,914 |
| 72h | B | 34,825,718 | 33,309,312 | 29,991,355 | 28,917,995 | 28,184,071 | 32,004,826 |
| 144h | A | 26,521,109 | 32,068,870 | 31,701,845 | 28,877,883 | 31,308,395 | 34,820,920 |
| 144h | B | 36,076,103 | 35,384,106 | 33,676,271 | 31,871,680 | 35,266,910 | 37,216,213 |

| Sample | Replicate | Map Rates |  |  |  |  |  |
| --- | --- | --- | --- | --- | --- | --- | --- |
|  |  | IR29 | Pokkali | FL478 | FL510 | FL499 | FL454 |
| 0h | A | 80.54% | 73.60% | 83.83% | 92.98% | 93.03% | 91.33% |
| 0h | B | 78.62% | 71.49% | 81.77% | 90.12% | 90.12% | 88.73% |
| 24h | A | 68.20% | 82.29% | 63.37% | 93.90% | 93.72% | 86.42% |
| 24h | B | 67.39% | 80.88% | 62.66% | 92.03% | 92.32% | 84.28% |
| 48h | A | 94.33% | 94.60% | 94.57% | 94.30% | 94.54% | 94.50% |
| 48h | B | 92.69% | 93.38% | 93.31% | 92.76% | 93.39% | 92.99% |
| 72h | A | 90.92% | 94.07% | 94.29% | 94.24% | 94.06% | 93.77% |
| 72h | B | 89.27% | 92.39% | 93.08% | 93.11% | 92.78% | 92.01% |
| 144h | A | 90.69% | 93.14% | 91.68% | 93.97% | 94.49% | 93.14% |
| 144h | B | 89.44% | 91.81% | 90.52% | 93.10% | 93.56% | 91.85% |

